# FoxP3 associates with enhancer-promoter loops to regulate Treg-specific gene expression

**DOI:** 10.1101/2021.11.12.468430

**Authors:** Ricardo N. Ramirez, Kaitavjeet Chowdhary, Juliette Leon, Diane Mathis, Christophe Benoist

## Abstract

Gene expression programs are specified by higher-order chromatin structure and enhancer-promoter loops (EPL). T regulatory cells (Treg) identity is dominantly specified by the transcription factor FoxP3, whose mechanism of action is unclear. We applied proximity-ligation with chromatin immunoprecipitation (HiChIP) in Treg and closely related conventional CD4+ T cells (Tconv). EPL identified by H3K27Ac HiChIP showed a range of connection intensity, with some super-connected genes. TF-specific HiChIP showed that FoxP3 interacts with EPLs at a large number of genes, including some not differentially expressed in Treg vs Tconv, but enriched at the core Treg signature loci that it upregulates. FoxP3 association correlates with heightened H3H27Ac looping, as ascertained by analysis of FoxP3-deficient Treg-like cells. There was marked asymmetry in the loci where FoxP3 associated at the enhancer- or the promoter-side of EPLs, with enrichment for different transcriptional cofactors. FoxP3 EPL intensity distinguished gene clusters identified by single-cell ATAC-seq as co-varying between individual Tregs, supporting a direct transactivation model for FoxP3 in determining Treg identity.

**One Sentence Summary:** FoxP3 is associated with enhancer-promoter loops in Treg cells, and correlates with heightened enhancer-promoter cross-talk

## INTRODUCTION

Looping is the solution that allows the packaging of the gigantic strands of chromosomal DNA into tight nuclear spaces, while ensuring the spatial organization and connectivity essential for the orderly deployment of their coding capacity (*1–3*). Some of the resulting topologies are very large, on the >megabase scale, such as the higher-order compartment domains, and primarily relate to overall organization. More directly associated with the unfolding of the genome’s coding potential are the enhancer-promoter loops (EPL) that bring distal enhancers into close proximity to transcriptional startsites (TSS) in the 3D space, enabling their interactions to regulate transcriptional programs (*4–6*). Rewiring of EPLs occurs during differentiation in several mammalian systems, ranging from early differentiation to olfactory receptor allelic choice (*7–10*). It has been proposed (*5*) that programmatic shifts associated with differentiation involve new enhancer-promoter connectivity, while the fast responses to stimuli or growth factors exploit pre-existing contacts, by modifying the activity of associated proteins (*11–13*). This dichotomy may not be as sharp as stated, and it is unclear how much EPL rewiring exists between closely related cells. Loops are dynamic structures, and the burst-like nature of transcriptional output may correspond to transient enhancer-promoter interactions (*14*). Finally, how sequence-specific transcription factors (TFs) coopt the EPL landscape remains mostly conjectural. Intuitively, fast-acting TFs whose mechanism of action relies on nuclear translocation (e.g. NF-AT) or activation at the cell membrane (e.g. STATs) likely operate on pre-existing EPLs. Other TFs might, instead, provoke the formation of new EPLs by dimerization (*15*) or macromolecular complex formation. A corollary question is whether an EPL is intrinsically sufficient for transactivation, or only a framework onto which activation mechanisms latch (*4, 5*). The regulatory function of TFs operating via EPLs has been demonstrated for both cell-specific and general TFs (e.g. Klf4 and Yy1, respectively) (*16, 17*).

T regulatory (Treg) cells are a branch of CD4+ T cells, dominant negative regulators of many facets of the immune system (*18, 19*). They also control extra-immunologic consequences of inflammation in several organs (*20*). Consistent with these pleiotropic functions, several phenotypic variants of FoxP3+ Tregs exist, with tuned transcriptomes that adapt to their tissue localization and effector functions (*21*). FoxP3, a member of the forkhead/winged-helix family of TFs, is quasi-exclusively expressed in Treg cells, and plays a central role in determining their identity and function, and their characteristic transcriptional signature. It is, however, neither completely necessary nor sufficient for Treg determinism (*22–25*), and requires synergistic action from a number of transcriptional cofactors with which it interacts (*26–31*). FoxP3 is not a pioneer factor, as many of the enhancers and open chromatin regions (OCR) to which it binds are already accessible in the T cell lineage prior to FoxP3 expression (*32, 33*), although a smaller group of FoxP3-binding OCRs are activated contemporaneously with Treg differentiation.

While FoxP3 can synergize with several sequence-specific TFs and with chromatin modifiers, its true mechanism of action remains poorly understood. There is debate as to it being an activator or a repressor, and even whether it activates Treg-specific transcription directly or by tuning intermediates. Some have argued that it functions primarily as a transcriptional repressor (*26, 27, 34-36*). The repression of cytokines produced upon activation by conventional CD4+ T cells (Tconv), and in particular interleukin (IL)-2, is a well-established function of FoxP3. On the other hand, results from several experimental systems suggest that FoxP3 also behaves as an activator for many of its transcriptional targets (*31, 37–40*), and that this is its principal mode of action. The “cofactor model” posits that this functional dichotomy depends on the identity of the cofactor(s) with which it interacts in controlling specific targets (*27, 31, 38*). Super-resolution microscopy does show FoxP3 associated with activating or repressive co-factors in different “transcriptional hubs” in the nucleus of the same Treg cell (*31*).

An important missing piece of the FoxP3 puzzle is an understanding of its interactions with nuclear loop structures, in particular enhancer-promoter connectivity. Chromatin immunoprecipitation (ChIP-seq) data show FoxP3 binding in the vicinity of TSS and at distant enhancers (*31, 32, 41*), but how FoxP3 inter-relates with EP connectivity is unknown. Does FoxP3 enhance, or disrupt, the formation of specific EPLs in Tregs? Does it dock onto existing loop structures to nucleate regulatory (activating or inhibitory) complexes? Here, we tackle this question by using chromatin proximity ligation with immunoprecipitation (HiChIP), first with anti- H3K27Ac antibody to establish the comparative landscape of EPLs in primary Treg and Tconv cells, two closely related cell-types. We then map FoxP3 molecules onto Treg EPLs. Directly relevant to the mystery of FoxP3 mechanics, the results show that FoxP3-bearing EPLs correspond to strong H3K27Ac loops genomewide, and positively associate with differential transcription between Tconv and Treg cells, consistent with action as a direct transcriptional activator.

## RESULTS

### Mapping the enhancer-promoter connectome in Treg and Tconv cells

To understand how the short- and long-range enhancer-promoter contacts mediate the cell type-specific expression patterns among mouse CD4+ T cells, we first used the “HiChIP” method (*9*), which identifies chromatin loops associated with a histone mark or a particular TF by combining proximity ligation of DNA strands with immunoprecipitation, quantitated by high-throughput sequencing. Antibodies against H3K27Ac, the histone modification typical of active enhancers, focused the analysis on loops connecting active enhancers. We analyzed primary mouse Treg and Tconv cells directly purified *ex vivo* to avoid imprints from cell culture, aiming for deep libraries to support robust quantitative analysis (200-300 M reads/sample on average). This required sizeable numbers of cells, achieved by magnetic purification from *Foxp3-Thy1.1* reporter mice. In line with current practice (*42, 43*), we quantitated EPL signals as reads that connect the promoter region (+/-500 bp around the TSS) to 5 kb bins tiled +/-250 kb from the gene’s TSS. To correct for proximity-biased background in HiChIP data, we compared the EPL signals to a null distribution generated from the interactions of 1000 random intragenic positions, which allowed estimates of significance corrected for relative distance to the TSS (similar to (*42–44*)). Signals were conserved among biological replicates (fig. S1A). A control HiChIP with an irrelevant isotype-matched antibody showed that the vast majority of H3K27Ac+ EPLs were indeed specific (p <0.0001, Fig. S1B).

Across both Treg and Tconv, 115,538 significant EPLs were identified (FDR 0.05) among 9,058 genes selected for reliable mRNA expression in Treg or Tconv cells (Table S1), as illustrated for *Lrrc32* in Fig1A. The median H3K27Ac EPL size was 49.5 kb (S1C), concordant with EPL size estimates in other mammalian cells (*7, 9*). Only 0.5% of H3K27Ac EPLs reached further than 1 Mb from the TSS, indicating that such very-long-range interactions can occur, but are the exception. Cumulative EPL intensity for each gene was only marginally related to its mRNA levels (fig. S1D), fitting with recent reports (*5, 45*).

A gene’s cumulative EPL intensity showed a linear relationship to its number of EPLs (Fig. 1B). At first blush this is mathematically obvious, but on further thought it implies that an active promoter does not simply allocate a fixed interaction potential by alternating between its potential enhancers, but that each enhancer has a given probability of interaction, and that these are cumulated for each gene (plausibly by co-recruitment into a transcriptional hub). This continuity suggests that the regulatory crosstalk of enhancers that loop to the same promoter operates through additive modalities (*46*).

**Figure 1.**
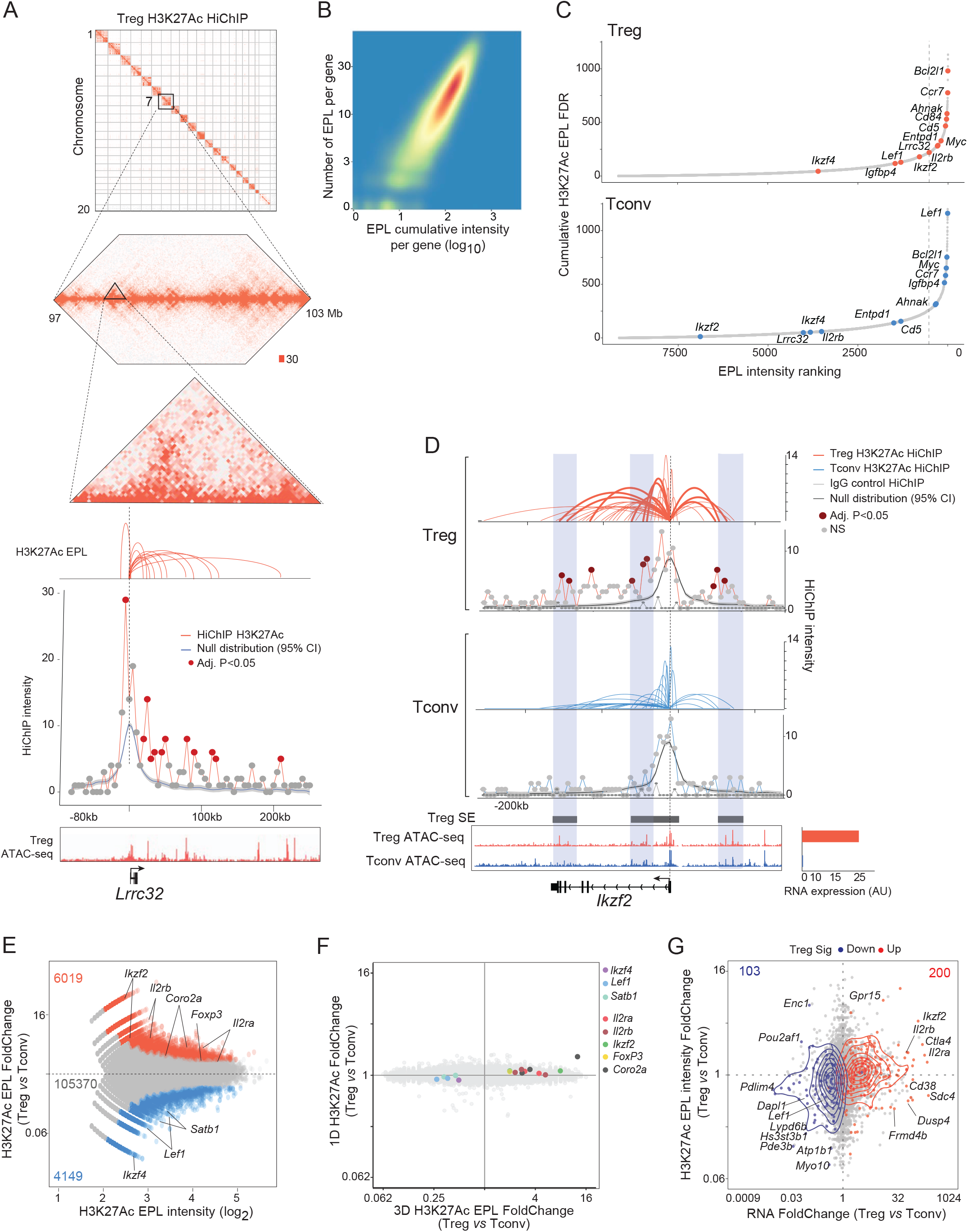
T-cell specific enhancer-promoter loops (EPLs) sync with the transcriptional regulation of the Treg gene program. A. Profiling the genome-wide enhancer-promoter architectures of mouse regulatory T cells *in vivo* with H3K27Ac HiChIP. A focused view of the *Lrrc32* locus shows a null distribution based on random proximity-biased ligations (95% CI) and those called as significant H3K27Ac HiChIP enhancer-promoter loops (FDR 5%) in Tregs. B. Scatter density plot of Treg H3K27Ac HiChIP EPLs and the relative cumulative intensities (Pearson r=0.78, log10). C. Ranking plots of super-interactive promoters determined for H3K27Ac Treg (top, n=523 genes) and Tconv (bottom, n=520 genes) using the summed cumulative H3K27Ac EPL FDRs respectively. D. T cell-specific enhancer-promoter configurations of the *Ikzf2* locus in Treg (top) and Tconv (bottom) cells. Differential EPLs are considered as >2 fold change and adj.P<0.05. IgG control HiChIP and null distributions (95% CI) are also shown for Tregs and Tconvs. ATAC-seq genomic tracks and RNA expression for *Ikzf2* are shown for Tregs and Tconvs respectively. E. Fold change vs. mean intensity (MA) plot of differential H3K27Ac HiChIP EPLs between Treg (red) and Tconv (blue) cell-types. EPLs with differential intensity (adj.P<0.05, FC>2) are highlighted with numbers shown. F. Comparison of 3D H3K27Ac HiChIP EPL intensities (x-axis, fold change in Treg vs. Tconv) and 1D H3K27Ac HiChIP EPL intensities (y-axis, fold change in Treg vs. Tconv). G. Comparison of RNA expression (x-axis, fold change in Treg vs. Tconv) and cumulative H3K27Ac EPL intensities (y-axis, fold change in Treg vs. Tconv). Genes are colored as Up- (red) or Down- (blue) Treg signature membership.

Some genes had far more EPL connectivity than others, over a 10-fold range (Fig. 1C, Table S2). Such a spread was observed in non-immunologic lineages, where some authors proposed to distinguish a class of “Super Interactive Promoters” (*47, 48*). Here, such super-connected genes (523 in Treg, 520 in Tconv) include several Treg and Tconv hallmarks like *Entpd1, Lef1, Ccr7, Cd5,* or *Lrrc32*. These super-interactive genes tend to be associated with “Super Enhancer” elements: of the 65 super-enhancers nominated in Tregs (*41*), 13 included super-interactive promoters (chisq p <0.01).

Differences between Treg and Tconv were readily identified, as exemplified by several prototypical Treg signature transcripts. Several EPLs were observed at the *Ikzf2* promoter in Tregs, but none significantly above background in Tconv (Fig. 1D). ATAC-seq accessibility patterns were essentially identical in Treg and Tconv at these positions, however, suggesting that the locus is poised but not connected in Tconv (which do express *Ikzf2* upon activation). Reciprocal Treg or Tconv patterns were observed for Treg signature genes *Il2rb* and *Pde3b* (fig. S1E). Genome-wide, we identified 10,168 (9% of total EPLs) differentially active EPLs (at fold-change >2, p <0.05; Fig. 1E, fig. S1F). Signal intensities in HiChIP data integrate true loop frequency and stability (“3D signal”) with the total abundance of the mark targeted by the immunoprecipitation (“1D signal”) (*9*). Total EPL signal in HiChIP data could thus appear to change, even if there was no actual increase in loop abundance *per se*. Analysis of the variation of total (1D) H3K27Ac signal showed that the Treg/Tconv differences in EPL intensity did represent increased looping, as they could not be accounted for by simple variation in H3K27Ac deposition, which varied very modestly (Fig. 1F). This degree of EPL variation between CD4+ T subsets is generally more extensive than the disparity observed by chromatin accessibility (less than 1% difference in accessibility) (*32*). Some genes that prototypically distinguish Treg and Tconv show differential EPLs (*Lef1, Satb1, Il2ra, Foxp3,* fig. S1G), consistent with observations in human T cells (*9*). Accordingly, pairing of gene expression and cumulative EPL intensity revealed a positive relationship between differential EPL intensity and mRNA levels between the two cell-types (Fig. 1F).

### FoxP3 preferentially targets hyperactive enhancer-promoter loops

Having established the overall landscape of EPLs in Treg and Tconv cells, we performed HiChIP with anti-FoxP3 antibody to identify enhancer-promoter interactions that involve FoxP3, for clues to understand how it performs its central role in shaping Treg identity. Data were generated in well-correlated biological duplicates (fig. S2A), with correction for proximity ligation and an irrelevant IgG control to ensure specificity (fig. S2B-C). To help identify the most reliable FoxP3 EPLs, we leveraged existing FoxP3 ChIP-seq data (*32, 41*) (fig. S2D), and filtered putative FoxP3 EPLs against this input. (fig. S2E). We thus identified 13,681 robust FoxP3-decorated EPLs, of ∼ 20 kb median length (fig. S2F, 2A). FoxP3 EPLs connected enhancers to the promoters of a surprisingly large number of genes (4, 797), confirming that FoxP3 relates to far more loci than just the ∼500 genes of the Treg signature. Here again, a subset of genes showed super- connectivity (Fig. 2B), enriched for T cell-specific ontologies (T activation p=10^-12^, cytokine signaling p=10^-11^) and Treg signature genes (*Il2ra, Ikzf2, Entpd1*). As illustrated for *Entpd1* and *Il2ra*, prototypical Treg-Up signature genes, abundant FoxP3-decorated EPLs corresponded to FoxP3-binding enhancers identified by ChIP-seq (Fig. 2A, S2G) and overlapped with a subset of the H3K27Ac EPLs. At both the *Lef1* and*Tcf7* loci, which are under-expressed in Tregs, FoxP3 EPLs also aligned with the positions of H3K27Ac EPLs, the latter being far less intense in Treg than in Tconv cells, consistent with an inhibitory impact of FoxP3 (Fig. 2C, S2H). At the *Il2* locus, two H3K27Ac EPLs were observed in Tconv (fig. S2I), but no loops whatsoever were detected in Treg, whether in association with H3K27Ac or with FoxP3, suggesting that the locus is closed into heterochromatin independently of FoxP3.

**Figure 2.**
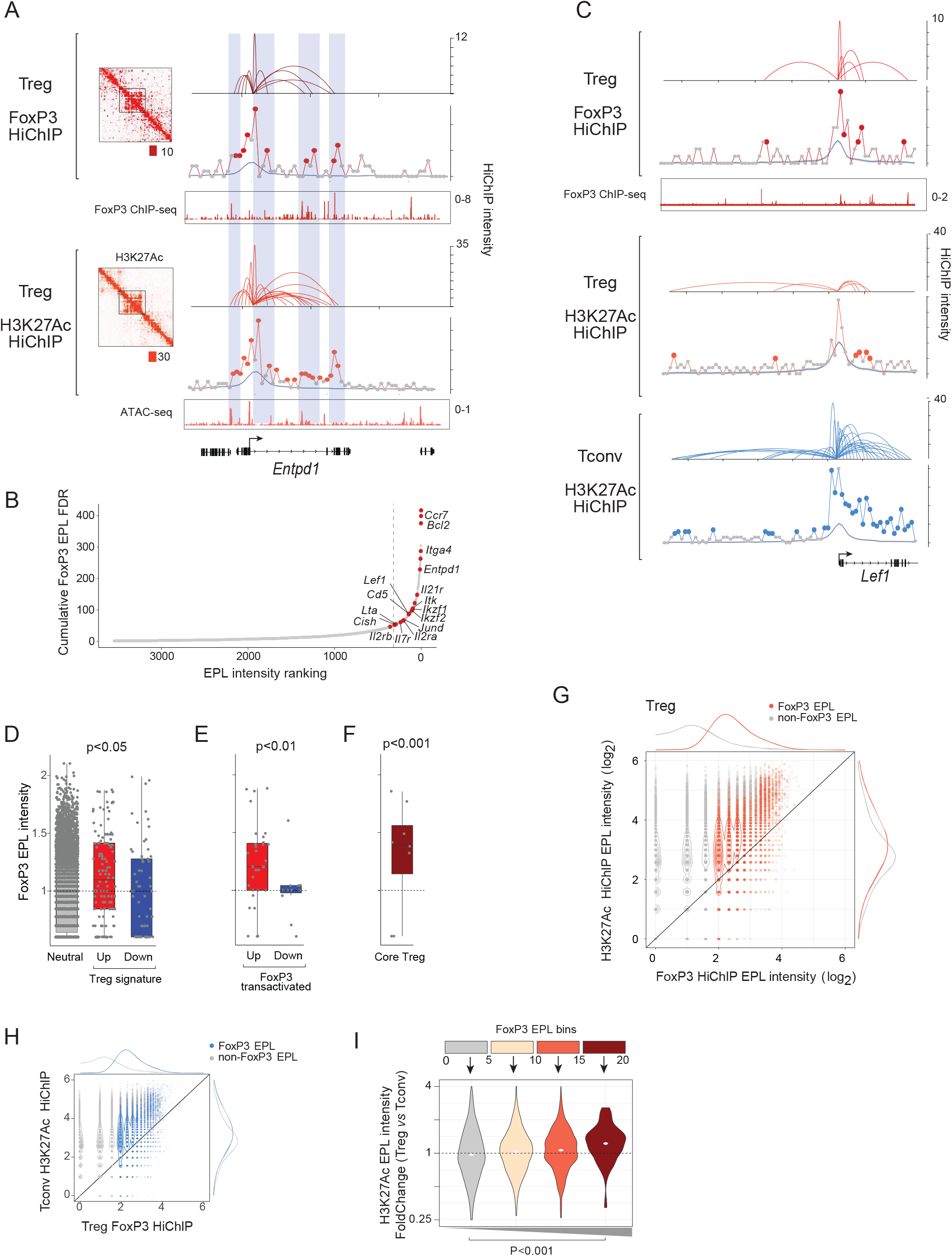
Genome-wide Treg FoxP3 enhancer-promoter looping generally associates with H3K27Ac-marked architectures. A. Treg FoxP3 (top) and Treg H3K27Ac (bottom) enhancer-promoter loops shown for the target gene *Entpd1.* Highlighted areas emphasize the agreement between FoxP3 and H3K27Ac EPLs. B. Rank plot of the FoxP3-specific super-interactive promoters (n=320 genes). C. FoxP3 (top), Treg H3K27Ac (middle), and Tconv H3K27Ac (bottom) enhancer-promoter loops for the *Lef1* locus. D-F. Boxplots of FoxP3 cumulative EPL intensities for the Treg (D), FoxP3 transactivated (E) and Core signatures (F). Significance was determined for Treg Up (p<0.05), FoxP3 transactivated Up (p<0.001), and Core Treg (p<0.01) genesets using a t-test. H. Scatter plot of FoxP3 HiChIP EPL intensities (x-axis, log2) and Treg H3K27Ac HiChIP EPL intensities (y-axis, log2). FoxP3-specific EPLs are highlighted. I. Scatter plot of FoxP3 HiChIP EPL intensities (x-axis, log2) and Tconv H3K27Ac HiChIP EPL intensities (y-axis, log2). FoxP3-specific EPLs are highlighted. J. Fold change distributions comparing H3K27Ac HiChIP EPL intensities (y-axis, Treg vs. Tconv) for each FoxP3 EPL bin (x-axis). Significance was determined using a t-test (p<0.001).

We then analyzed the distribution of FoxP3 EPLs, beyond these examples, in genesets that reflect differential expression in Treg cells and FoxP3-dependence. Classic “Treg signature genes”, which are known to have FoxP3-dependent and independent components (*39, 49*), showed a modest but significant increase in cumulative FoxP3 EPL intensity compared to expression-matched “neutral” genes (Fig. 2D). This enrichment was accentuated in a set of FoxP3-dependent genes that are up-regulated after short-term FoxP3 transfection (*31*) (Fig. 2E). No such enrichment was seen for down-regulated targets (Fig. 2D,E). The strongest enrichment in FoxP3-associated EPLs was seen for genes that constitute a core signature of FoxP3- dependent genes identified in FOXP3-deficient mice and patients (*25*), Fig. 2F). These results are consistent with the notion that transcriptional upregulation by FoxP3 is directly achieved through enhancer-promoter connections at up-regulated loci.

Overall, plotting the relationship between FoxP3 and H3K27Ac EPLs showed that FoxP3 EPLs correspond to loops with the highest level of H3K27Ac-associated signals (Fig. 2G), suggesting that FoxP3 associates with the most active chromatin. To ensure that this observation was not an experimental artefact, we performed HiChIP for the Yy1 TF, which is considered a structural regulator broadly involved in enhancer-promoter topologies (*17, 50*). In Tregs, Yy1 represses *Foxp3* (*51*), and inhibits transactivation by FoxP3 protein (*31, 51*). The 12,426 Yy1 EPLs thus identified were also enriched in H3K27Ac HiChIP intensity when compared to non-Yy1 chromatin interactions (p<0.05), similar to FoxP3, and consistent with observations in stem cells (*17*). But FoxP3 and Yy1 EPLs showed differential EPL patterns (fig. S4A,B), most strikingly at *Il2* (fig. S2H). In a combined set of FoxP3/or/Yy1 EPLs (n=21,341), 54.3% were differentially represented (>2-fold difference in intensity). The genes associated with FoxP3 or Yy1 EPLs belonged to different functional classes: Yy1 EPLs were enriched in “General transcription” or “Cell Protein Catabolism” ontologies, while FoxP3-associated EPLs belonged to more immune-specific pathways (“Adaptive immune system”, “MHC-I presentation”; fig. S4C).

Examination in Tconv of those H3K27Ac EPLs that bind FoxP3 in Tregs revealed that most of them also had high intensity in Tconv (Fig. 2H). Thus, FoxP3 binds to enhancer-promoter loops that are very active in CD4+ T cells in general, and not solely to Treg-specific ones, consistent with the notion that it homes to chromatin locales that are ubiquitously open in T cells (*32, 33*). If FoxP3 locates to Treg EPLs with the highest degree of activation, how does its presence affect them? To address this question most finely, we compared the H3K27Ac EPL intensities of the EPLs in Tregs, where FoxP3 is bound, versus in Tconv where FoxP3 is absent, across bins of FoxP3 EPL intensity. Increasing FoxP3 corresponded to increased H3K27Ac EPL signal at those EPLs in Tregs relative to Tconv (Fig. 2I), suggesting that FoxP3 may exert a facilitating influence on enhancer-promoter loops.

### Direct tuning of FoxP3-dependent transcriptional programs through FoxP3 enhancer-promoter loops

The results are compatible with, but do not prove, the notion that transcriptional activation by FoxP3 is actuated directly through enhancer-promoter connections around its target genes. However, this notion is at odds with the recent claim that FoxP3 shapes Treg identity indirectly, by regulating the abundance of intermediary TFs like TCF1 (encoded by *Tcf7)* (*52*). While we did observe FoxP3-decorated EPLs across the *Tcf7* locus (fig. S2H), the sweeping conclusion that indirect regulation shapes Treg transcriptional identity is likely an overstatement, as discussed in detail in the Supplementary Note. *Some* indirect control probably occurs (since FoxP3 controls several other TFs), but a sizeable portion of the FoxP3-dependent transcriptome is likely to be directly controlled by FoxP3, including key functional transcripts. It was important, however, to verify that the FoxP3-decorated EPL structures reported here are functionally relevant. To this end, we performed H3K27Ac HiChIP on Treg-like cells sorted from a novel FoxP3-deficient line (hereafter KO-GFP, constructed from the classic FoxP3-ires-GFP reporter mouse (*54*) by a CRISPR-engineered frameshift mutation that eliminates the Forkhead domain resulting in a full *scurfy*-like Treg deficiency phenotype – see Methods). As other FoxP3-deficient mice (see (*25*) for refs), KO-GFP mice contain a sizeable proportion of FoxP3-null Treg-like cells, identifiable by activity of the GFP reporter appended to the disarmed *Foxp3* locus. Heterozygous females with one copy of this locus are protected from disease by the WT *Foxp3* allele (provided by a *Foxp3- Thy1.1* reporter chromosome to facilitate identification). Genomic comparison to cells in which the intact Foxp3-GFP allele (hereafter WT-GFP) is active enables very precise determination of the effect of the FoxP3 deficiency in Treg-like cells, identified by the same reporter configuration. We sorted WT-GFP and KO-GFP Tregs from half-sib heterozygous females (Fig. 3A) and performed H3K27Ac HiChIP to evaluate FoxP3’s effects on enhancer-promoter looping.

**Figure 3.**
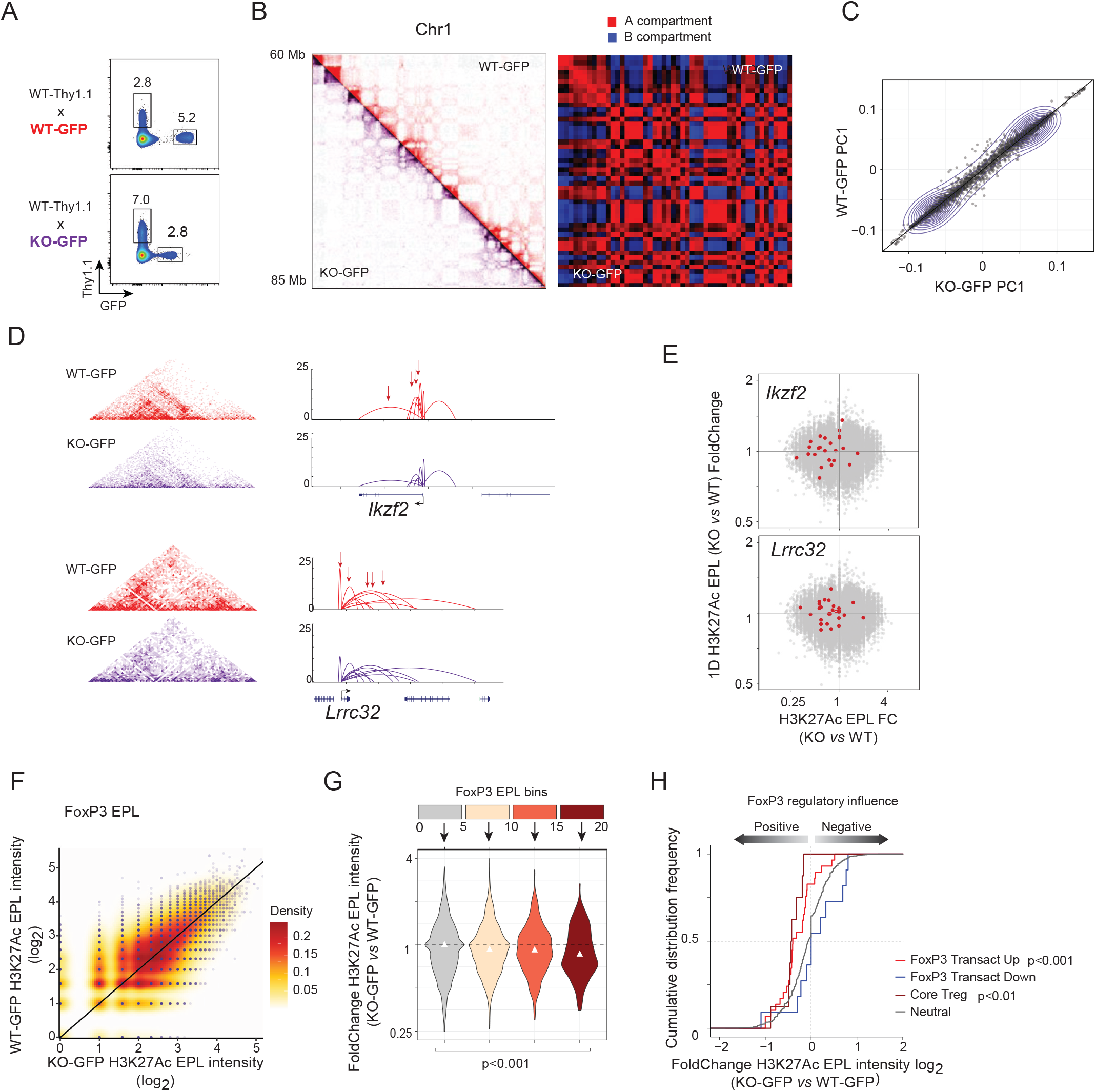
FoxP3 enhancer-promoter looping directly modulates FoxP3-dependent transcriptional program. A. FACS plots for WT-GFP (top) and KO-GFP (bottom) sorted Treg populations (gated on the CD4+TCRB+ population). B. WT-GFP and KO-GFP intensity and chromatin compartment H3K27Ac HiChIP maps. C. Genome-wide scatter plot of H3K27Ac HiChIP genome compartment scores (Pearson 0.98). D. WT-GFP (red) and KO-GFP (purple) H3K27Ac HiChIP intensity maps (left) and EPLs (right) of FoxP3 targets *Ikzf2* and *Lrrc32*. Arrows indicate EPLs FC>1.5 and P<0.05). E. Comparison of 3D H3K27Ac HiChIP EPL intensities (x-axis, fold change in KO-GFP vs. WT-GFP) and 1D H3K27Ac HiChIP EPL intensities (y-axis, fold change in KO-GFP vs. WT-GFP). *Ikzf2* and *Lrrc32* specific H3K27Ac EPLs are indicated. F. Scatter plot of WT-GFP and KO-GFP H3K27Ac HiChIP EPL intensities for FoxP3- specific EPLs. G. Fold change distributions comparing H3K27Ac HiChIP EPL intensities (y-axis, KO-GFP vs. WT-GFP) for each FoxP3 EPL bin (x-axis). Significance was determined using a t- test (p<0.01). H. Cumulative distribution frequency plot comparing KO-GFP and WT-GFP H3K27Ac EPL intensities for FoxP3 Transactivated (Up, p<0.001) and Core Treg (p<0.01) signatures. Significance was determined using a two-sided KS.test.

Lieberman-Aiden et al showed that HiC and HiChIP data, when analyzed by Principal Component Analysis, reveals spatial segregation of open and closed chromatin that forms two genome-wide compartments (*55*). By this measure, FoxP3-deficiency did not affect nuclear compartments on the macro scale, with no change in heterochromatin compartments identified by Principal Component Analysis (Fig. 3B,C). However, loci with rich FoxP3 EPLs, like *Ikzf2* and *Lrrc32,* showed significantly reduced H3K27Ac-decorated loops in FoxP3 KO Tregs (arrows in Fig. 3D, p<0.05); these differences were reproducible in biological replicates (fig. S3F). As above, this reduction in EPL intensity in KO-GFP Tregs denoted a true loss of connectivity, not merely lower H3K27Ac decoration (Fig. 3E): the drop in H3K27Ac EPL intensity (x-axis) was not accompanied by a drop in total “1D” H3K27Ac signal at these loci. Conversely, FoxP3 down- regulated targets *Lef1* and *Add3* showed increased H3K27Ac EPL signal in KO-GFP Tregs (fig. S3G). Comparing genomewide H3K27Ac EPL abundance in WT-GFP and KO-GFP Treg-like cells showed a general concordance, with only a light bulge for the most highly represented EPLs (Fig. 3F). This deviation was confirmed by plotting the KO-GFP/WT-GFP ratio of H3K27Ac EPL intensity across FoxP3 EPL intensity bins (Fig. 3G, p<0.01), showing a clear reduction in signal resulting from the lack of FoxP3 in those EPLs with highest FoxP3 EPL intensity. The conclusion that FoxP3 directly shapes chromatin connectivity, and its transcriptional relevance, were confirmed by the cumulative density plot of KO/WT ratios for FoxP3 target genes defined as above (from direct transactivation studies or the core Treg set - Fig. 3H). Enhancer-promoter loops in FoxP3-upregulated genes were more sensitive than the norm to the absence of FoxP3 (and the converse was true of FoxP3-repressed loci). These results support FoxP3’s association to epigenetically active loops, though which it directly influences the Treg-specific gene program.

### Two different categories of FoxP3-dependent genes

These results show that, as a group, transcripts whose expression is upregulated by FoxP3 are enriched in FoxP3-adorned EPLs, but this characteristic does not necessarily apply to all such genes. To further discriminate between loci with FoxP3-enhanced expression in Tregs, we compared accessibility in single-cells: similarly regulated genes tend to have correlated activity across individual cells, and correlation analysis can uncover groups of genes with similar regulatory logic (*56, 57*). We used single-cell ATAC-seq (scATAC-seq), as chromatin accessibility is more stable than mRNA levels (*57–59*), exploiting a dataset generated for another study (KC, in preparation), with high-quality scATAC-seq profiles for 5,810 splenic Tregs, including rTreg and aTreg types (Fig. 4A). We first defined a FoxP3-dependent geneset from the expression data of van der Veeken at al (*52*), by contrasting Treg-like cells from heterozygous female mice in which FoxP3 deficient or proficient X-chromosomes were active (fig. S4D; 377 upregulated genes in FoxP3 proficient vs deficient at FC>2 and p<0.01). We generated a gene accessibility score from the scATAC-seq data and correlated these scores across all Treg cells in the scATAC dataset. This gene-gene correlation network resolved into two predominant gene clusters (Fig. 4A), which exhibited a marked imbalance in FoxP3 EPL density (Fig. 4B,C; p<0.01, Table S3). Loci in Cluster1 had high FoxP3 EPL density (Fig. 4B-D), and included many key Treg transcripts (*Il2ra*, *Il2rb*, *Lrrc32, Izumo1r* and *Nrp1*), predominantly characteristic of rTregs; in contrast, genes of Cluster2 showed much lower FoxP3 EPL density, and corresponded mainly to transcripts enriched in aTregs (e.g. *Ccr8*, Fig. 4D). Accordingly, genes from Cluster1 showed higher binding to FoxP3 in published ChIPseq datasets (Fig. 4E), with H3K27Ac EPL intensity also biased for Cluster1 transcripts (Fig. 4F). Overall, Cluster1 genes tended to be expressed at somewhat higher levels than Cluster2 (median 341 tpm, 0.05-0.95 range [27-3576] vs 76, [26-1417), but genes in both clusters were up-regulated by FoxP3, and over-expressed in WT relative to FoxP3- deficient Treg-like cells (Fig. 4G).

**Figure 4.**
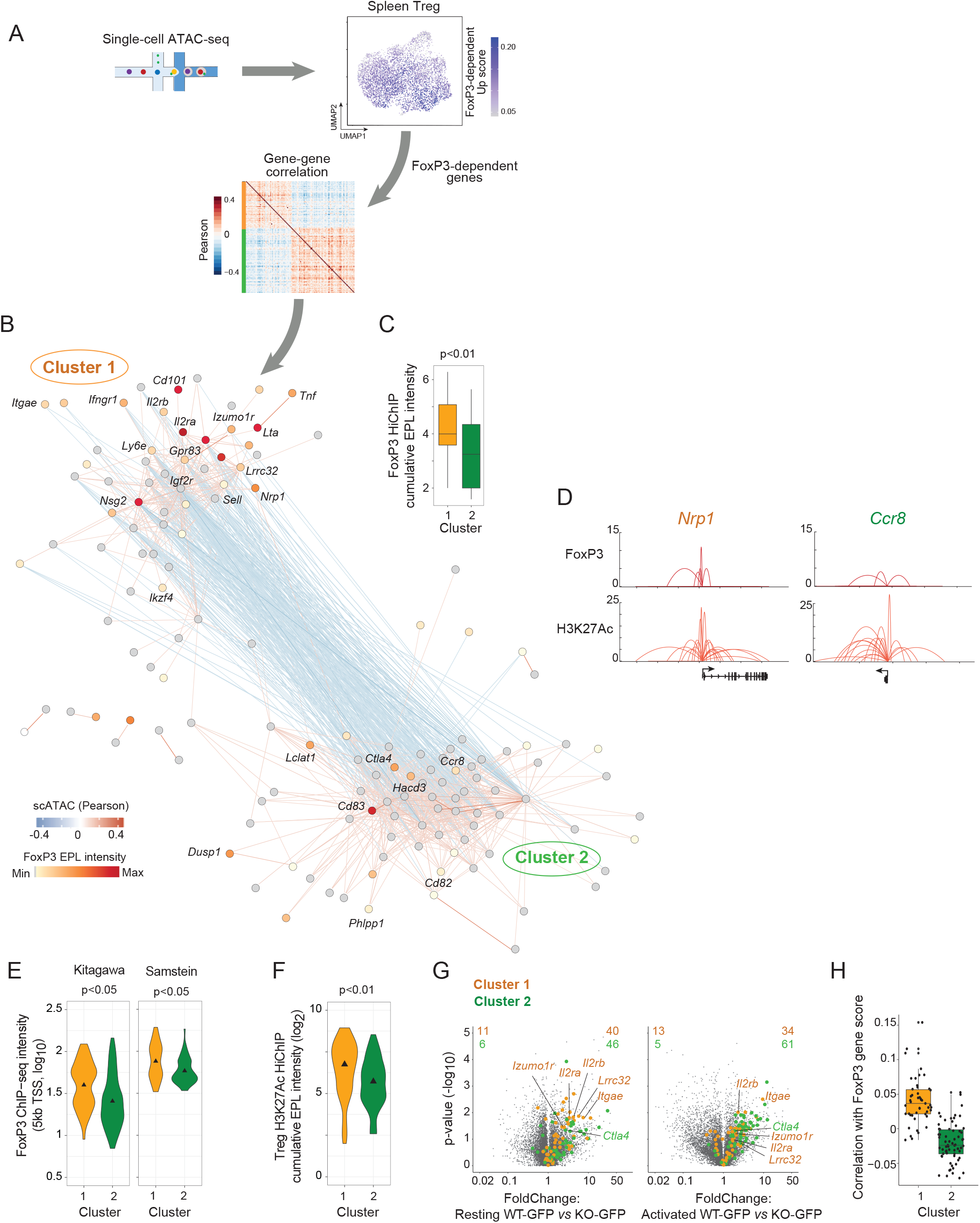
Parsing the FoxP3-dependent coregulated network with single-cell chromatin accessibility. A. Schema of splenic Treg profiling with single-cell ATAC-seq. UMAP of splenic Tregs representing the FoxP3-dependent ‘Up’ signature gene score and gene-gene correlation heatmap (Pearson). B. Gene scores were calculated from Treg scATAC-seq for the FoxP3-dependent signature and represented as a force-directed network. Gene-gene correlations (edges) and FoxP3 EPL cumulative intensities for each gene (node) in the network reflected by colored spectra. Indicated clusters were determined using both edge and node attributes. C. Box plot comparing FoxP3 HiChIP cumulative EPL intensities for Cluster1 (orange) and Cluster2 (green) genes. Significance was determined between groups using a t-test (p<0.01). D. FoxP3 and H3K27Ac HiChIP EPLs are shown for Cluster1 gene *Nrp1* and Cluster2 gene *Ccr8*. E. Violin plots of FoxP3 ChIP-seq intensities between Cluster1 and Cluster2 genesets. ChIP-seq signal was measured in a 5kb window around the TSS. F. Violin plots of Treg H3K27Ac cumulative EPL intensities between Cluster1 and Cluster2 genesets. G. Volcano plots comparing the transcript expression of resting and activated Tregs (WT- GFP vs KO-GFP). Gene clusters from (B) are highlighted. H. scATAC gene activity score correlations (Pearson) between FoxP3 and individual genes from Cluster1 and Cluster2 identified in (B).

We then asked, across all single-cell, how the expression of individual genes of Clusters 1 and 2 (approximated by the “gene accessibility score”) correlated with that of FoxP3. The rationale was that, while all these genes appear FoxP3 dependent (as defined by the “sledgehammer” effect of FoxP3 deletion) their instant expression may respond differently to fluctuations in FoxP3 between individual cells. Interestingly, Cluster 1 genes had some positive correlation with Foxp3 (Fig. 4H). In contrast, genes of Cluster2, that are largely devoid of FoxP3-decorated EPLs, lacked this correlation (was even slightly negative on average), This dichotomy might actually be what one would expect if Cluster1 were under direct and positive activation by FoxP3, while Cluster2 genes responded to indirect control by FoxP3. Thus, the partition of FoxP3 dependent genes into clusters independently marked by FoxP3 EPL density suggests that different regulatory mechanisms are at play in the different groups of FoxP3-dependent genes, with different levels of direct FoxP3 involvement.

### Distinct FoxP3 interaction at promoters and enhancers

FoxP3 binding across the Treg genome occurs at distal enhancers, as well as in proximal promoter regions (*32, 41*). It is unknown whether FoxP3 functions similarly in both locations. We assessed whether FoxP3 bound to the promoter or enhancer ends of the loops, based on the consensus ChIP-seq map described above. EPLs associated with FoxP3 on either promoter or enhancer sides, or both (Fig. 5A). This dichotomy was unrelated to the clusters of Fig. 4, and not merely due to sparse sampling or arbitrary thresholds, as FoxP3 binding was truly absent on the “other” side (Fig. 5B). Further, EPLs with FoxP3 on both sides were not a random occurrence, as they occurred more frequently than through chance association (chisq p<0.01). High chromatin accessibility and H3K27Ac signal intensity coincided with the side of the EPL bound by FoxP3 (fig. S5A), confirming FoxP3’s association with the most highly active chromatin. A clue as to the significance of FoxP3 placement relative to EPL ends came from Gene Ontology analysis of the corresponding genes. Generic programs linked to nucleic acid metabolism (e.g. Cell Cycle, DNA repair) dominated for genes with promoter-side FoxP3 EPL, while more immune-specific ontologies (Cytokine signaling and adaptive immune system) were enriched among enhancer- and dual-sided modes (Fig. 5C, Table S4). FoxP3-dependent loci belonging to Cluster1 described above were predominantly associated with enhancers (only 3 of 24 were of the promoter-side category).

**Figure 5.**
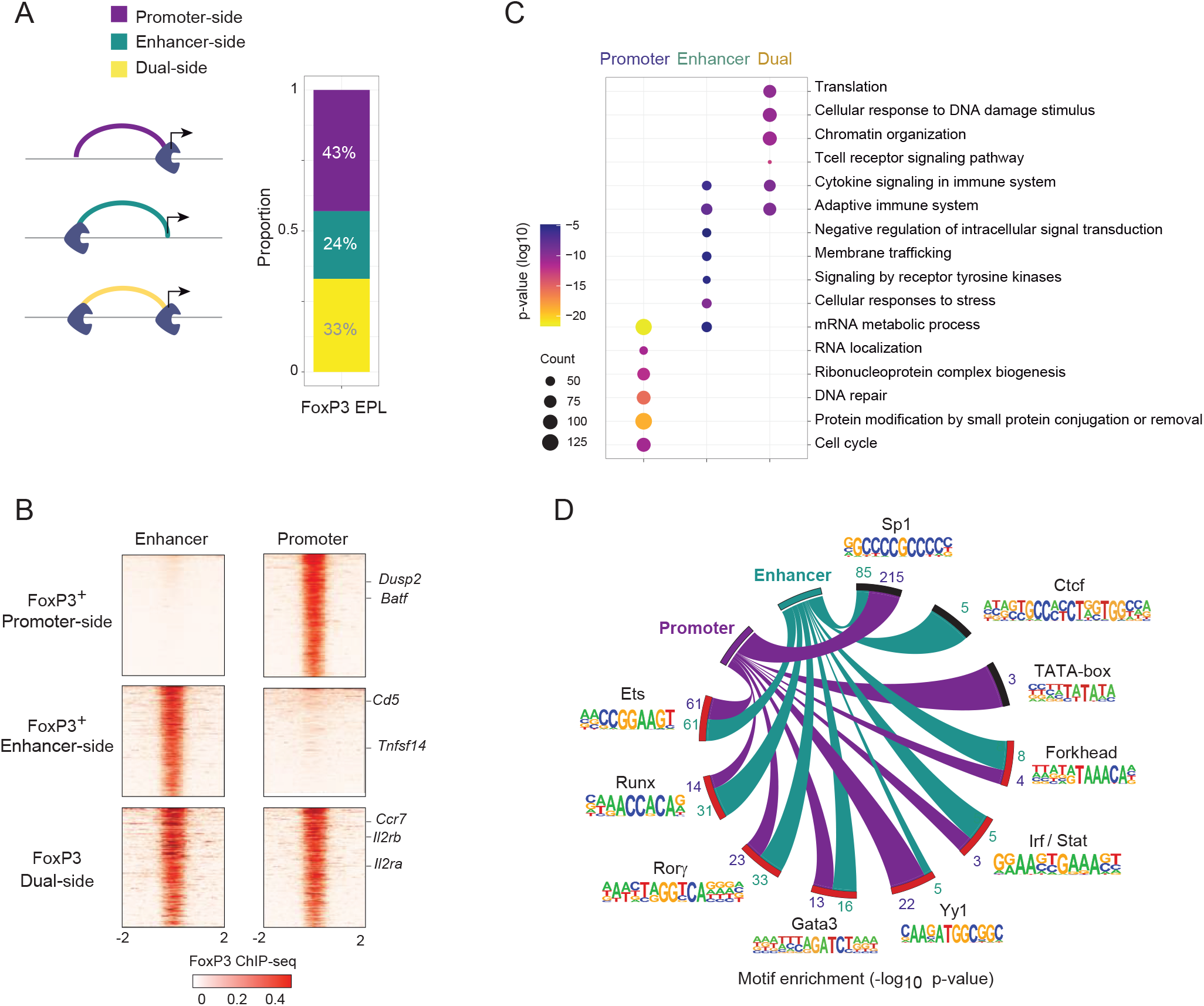
FoxP3 positioning at enhancer-promoter loops distinguishes gene and TF-motif components. A. FoxP3 HiChIP EPL proportions based on FoxP3 binding modes fixed on either the promoter, enhancer or dual sides of FoxP3 loops. B. FoxP3 ChIP-seq heatmaps centered on FoxP3 bound position within EPLs for promoter, enhancer and dual modes. C. Gene ontology enrichments based on FoxP3 HiChIP EPL mode. Size of circle indicates number of genes and color represents significance (p-value). D. Circos plot of TF motif enrichments for promoter and enhancer FoxP3 HiChIP EPLs. Arc thickness defines TF motif significance (-log10 p-value).

FoxP3 collaborates with other TFs, these interactions determining its transcriptional specificity (*26, 27, 31*), and their binding motifs are enriched in the vicinity of FoxP3 binding sites (*32, 41*). FoxP3 positioned at either the promoter or enhancer sides of EPLs appears to interact with distinct cofactors (Fig. 5D, S5B), as shown by motif enrichment analysis (using as background comparators all TSS and all non-TSS OCRs, respectively). TATA-box motifs were enriched at promoter positions, while Runx and Ctcf motifs were over-represented at enhancers, and motifs for SP1, Ets, Irf/Stat and the Nuclear Receptor family factors (Rorα/γ) were enriched on both sides. Together, these results delineate different aspects of FoxP3-mediated regulation, with distinctive transcription programs and TF cofactor preference across FoxP3 enhancer-promoter loops.

## DISCUSSION

To better understand the program controlled by the lineage-identifying factor FoxP3, we explored enhancer-promoter connectivity in true Treg and Tconv cells *ex vivo*, and how FoxP3 is associated with this architecture. The results lay out the landscape of enhancer elements that are connected to promoters of different genes in CD4+ T cells, and should thus serve as a roadmap to support cis-regulation studies in these important cells. Most importantly, and reinforcing the identity of FoxP3 as a positive transactivator, they establish how FoxP3 associates with a wide range of active EPLs, and positively potentiates their representation for a set of genes that are central to Treg cell physiology.

Previous models of enhancer regulation (*46, 60–62*) have hypothesized additive or even multiplicative enhancer activity. In the primary cells analyzed here, there was a significant but modest correlation between enhancer-promoter connectivity and mRNA levels in these two closely related CD4+ T cells. The absence of a linear relationship is consistent with the notion that simply establishing a connection between an enhancer and a promoter is not sufficient for transactivation, but that secondary events (recruitment of additional factors, post-translational modification of these cofactors) are necessary to functionally potentiate the connection. Cumulative EPL intensities for individual genes distributed as a very broad range, with a subset of highly connected T-cell genes, in line with recent work in neurons which identified a subset of “super-interactive promoters” (*48*). Such highly connected loci may be rationalized by the need for more complex transcriptional hubs for variegated regulation.

A key goal was to understand whether and how FoxP3 is involved with enhancer-promoter connectivity. Does it promote loop formation, or tune the transactivation potential of pre-established loops - the latter in keeping with the observation that, unlike other TFs that specify immunocyte lineages such as PU.1 or Pax5, FoxP3 opportunistically exploits already open chromatin and has limited “pioneer” activity - or does it have little influence, only acting indirectly, as suggested elsewhere (*52*). Generally, FoxP3 associated with the most epigenetically active chromatin, as did Yy1. Several lines of evidence established that, once associated with an EPL, FoxP3 mostly exerts an enhancing influence on enhancer-promoter connectivity, e.g. by stabilizing them: (i) H3K27Ac EPLs that are also associated with FoxP3 in Tregs are more intense in Tregs than they are in Tconvs; (ii) these same EPLs lose intensity when FoxP3 is genetically inactivated; in other words, when FoxP3 is absent, these enhancers are less connected to their cognate promoters. The functional relevance of these effects is established by the observation that these enhanced connections are found for genes whose expression is known to be positively transactivated by FoxP3.

Different generations of chromatin immunoprecipitation (*31, 32, 41, 52, 63, 64*) showed that FoxP3 associates with many genomic regions that are nowhere near the Treg-specific loci. In this, FoxP3 is not unlike many other TFs (*65*). Our results now show that these apparently inactive locations actually correspond to true enhancer-promoter connections, even at loci that have no clear Treg relevance (38% of Treg-neutral genes have a FoxP3-decorated EPL). An interpretation consistent with the “cofactor model” is that these positions simply lack the appropriate cofactors or epigenetic modifying enzymes, or that FoxP3 or its cofactors are not suitably modified at these locations in order to actually influence transcription. FoxP3 may be truly inert at these locations, or it might provide buffering or stabilizing influence (*61, 62, 65*). At FoxP3- dependent loci that are over-expressed in Treg cells, high-intensity FoxP3 EPLs observed at many core signature genes, which strongly suggest a positive and direct transactivation, in contrast to the indirect model recently proposed (*52*). In contrast, an indirect modality for FoxP3- dependent repression is consistent with the dearth of FoxP3 EPLs at the *Il2* locus. But indirect repression by FoxP3 is not a general characteristic of Treg-underexpressed loci, as *Lef1* and *Tcf7* genes themselves show abundant FoxP3-associated EPLs, consistent with the notion that FoxP3 can act as a direct repressor when paired with the appropriate cofactors (*31*).

Finally, the dichotomy between loci at which FoxP3 associates on the enhancer or the promoter sides of EPLs implies a geographical variation of its direct regulatory functions. This demarcation parallels recent results in B cells where distinct multi-TF hubs bind to promoter versus distal enhancer sites (*66*). The enrichment analysis (Fig. 5D) suggests that the factors with which FoxP3 associates on the enhancer or the promoter sides are different, suggesting different modes of operation. It is also possible that this repartition is dynamic (per (*12, 13*)), and one could speculate for instance that, upon rapidly-acting cell triggering, FoxP3 already bound to the promoter region serves to further stabilize/potentiate an EPL through dimerization with another FoxP3 molecule recruited to the enhancer side.

In terms of how FoxP3 determines Treg identity, parsing the group of FoxP3-dependent genes by correlation analysis of chromatin accessibility across single Tregs distinguished two groups of genes, among those positively controlled by FoxP3. The larger cluster was associated with abundant FoxP3-decorated EPLs, indicative of direct and positive transactivation. The second, smaller, cluster grouped genes associated with markedly less FoxP3-bound enhancer-promoter loops. Interestingly, genes of this second cluster include genes that are over-expressed in aTreg relative to rTreg, and these might correspond to indirect control (*52*).

These results, and those from (*25*) and (*52*), suggest a multimodal set of actions through which FoxP3 helps determine Treg identity and stability: (i) direct control, both activating and repressive, of a range of target loci, in particular a core set of transcripts essential for Treg homeostasis; (ii) repression of TFs that otherwise favor the Tconv program (iii) stabilization of a Treg program that is not directly FoxP3-dependent.

## METHODS

### Mice

C57BL/6J (B6), B6.*Foxp3^Thy1.1^* (*67*) mice were obtained from the Jackson Laboratory and bred in the SPF facility at Harvard Medical School (HMS). All experimentation was performed following animal protocols approved by the HMS Institutional Animal Use and Care Committee (protocols IS00000054). FoxP3-deficient mice (B6.*Foxp3^fs327-gfp/Doi^*) were generated by CRISPR mutagenesis of the B6.*Foxp3^ires-GFPKuch^* line (*54*) as described elsewhere (Leon et al, in preparation). Briefly, the Foxp3-ires-GFP allele carries a 1 bp insertion in exon 11 (NM_054039.2), at the position encoding FoxP3 amino-acid 327, leading to a frameshift mutation that would produce a truncated protein terminating after 34 amino-acids, and thus lacking the essential carboxy-terminal forkhead DNA-binding domain (fig. S3A). Males carrying the mutant allele present with a typical *Scurfy*-like phenotype of FoxP3 deficiency, with a wasting syndrome starting at 12-15 days of age (failure to thrive, cachexia, extensive exfoliative dermatitis, tail necrosis, crusty eyelids and ears, marked lymphadenopathy and splenomegaly) leading to death around 30-35 days. - Heterozygous females are healthy and don’t show any clinical signs of inflammation. The germline mutation was maintained onto the B6 background. For the experiments reported here, females carrying either the normal *Foxp3^ires-GFPKuch^* or the deficient *Foxp3^fs327-gfp^* alleles were crossed to B6.*Foxp3^Thy1.1^* father(the FoxP3-Thy1.1 reporter was used to identify in the offspring normal Treg cells that express the WT balancing allele in these crosses. To improve the comparability between parallel fs327 and WT offsprings in these experiments, the *Foxp3^ires-GFPKuch^* and *Foxp3^fs327-gfp^* carrier mothers were co-housed and bred to the same B6.*Foxp3^Thy1.1^* father.

### T cell isolation

For analyses in FoxP3 proficient mice, splenic CD4 T cells from FoxP3-Thy1.1 hemizygous male mice were first enriched using CD4 untouched kit (Miltenyi), followed by secondary purification with anti-CD90.1 microbeads (Miltenyi) according to manufacturer’s instructions. Tconv (FoxP3- Thy1.1-Tcrb+CD4+CD45+CD19-CD8-CD11b-CD11c-) and Treg cells (FoxP3- Thy1.1+Tcrb+CD4+, CD45+CD19-CD8-CD11b-CD11c-) were sorted on a FACS-Aria to 96% and 94% purity, respectively. For experiments in FoxP3-deficient Tregs (Fig. 3), CD4^+^ T cells from spleen and peripheral lymph nodes from 6 week-old heterozygous females were first enriched as above by negative magnetic selection (Stem Cell), and Treg-like cells sorted as CD19- TCRβ+CD4+Thy1.1-GFP+ on a Moflo Astrios (Beckman Coulter).

### H3K27Ac, FoxP3, Yy1, and IgG HiChIP

HiChIP (*9*) was optimized and performed in biological replicates for all conditions. Briefly, 5 x 10^6^ Tregs were crosslinked for each FoxP3 HiChIP replicate, 2 x 10^6^ million cells for H3K27Ac and Yy1 HiChIPs respectively. Additionally, 5 x 10^6^ million Tregs and Tconvs were crosslinked to include a negative IgG isotype control HiChIP samples. Low input H3K27Ac HiChIP was performed on 2 x 10^5^ crosslinked WT-GFP and KO-GFP Tregs respectively. DNA sonication of fragments to 200-600 bp following MboI digestion was performed using the Covaris M220. Sonicated DNA was pre-cleared using Protein A magnetic beads followed by overnight chromatin immunoprecipitations with 3 ug H3K27Ac (Abcam ab4729), 5 ug FoxP3 (Abcam ab150743), 3 ug Yy1 (Abcam ab109237) and 5 ug rabbit IgG (Cell Signaling 2729S), in a final concentration 0.07% SDS. Libraries were prepared using Illumina Nextera DNA library preparation kit, size-selected using low-melt gel extraction and paired-end sequenced using NextSeq 500 Illumina platform.

### scATAC-seq of Tregs

Single cell ATAC-seq (scATAC-seq) data were generated as part of another study (KC, in preparation). Briefly, splenic regulatory T cells (CD4+, TCRβ+, FoxP3-IRES-GFP+) and T conventional cells (CD4+, TCRβ+, FoxP3-IRES-GFP-) were sorted from 6-8 week old B6 FoxP3- IRES-GFP mice into DMEM + 2% FCS, then resuspended in PBS + 0.04% BSA. Nuclei isolation, transposition, GEM generation, and library construction targeting capture of ∼12000 cells (75% Tregs, 25% Tconv) were carried out as detailed in the Chromium Next GEM Single Cell ATAC manual (10x Genomics). Libraries were pooled and sequenced on an Illumina NovaSeq to a final median depth of approximately 30,000 reads per cell (∼80% saturation).

### RNA-seq expression data

Treg and Tconv RNA-seq biological replicates were obtained from previous work (*33*). RNA-seq counts were used for all gene quantification. Genes were filtered based on replicated and cell-type expression; > 10 counts in both replicates, and in at least 1 condition.

### ChIP-seq data

ChIP-seq datasets (*41*); Runx1, FoxP3, Ets1, Smc1a (Cohesin), (*32*); FoxP3, were mapped to mm10 genome using bowtie2 (*68*). BigWig files were generated using deeptools2 (*69*).

### ATAC-seq data

Treg and Tconv ATAC-seq datasets were generated in previous work (*33*).

### scATAC-seq analysis

Fastq data were aligned to the mm10 reference genome and quantified per cell using Cell Ranger ATAC software (10x Genomics, v1.2). Peak by cell count matrices were generated by mapping reads to open chromatin regions previously defined by the ImmGen consortium (*33*). Data analysis was performed using Signac v1.1 (***70***). For quality control, only cells with at least 4000 fragments per cell, greater than 55 percent reads in peaks, TSS enrichment score greater than 2, nucleosome signal less than 10, and ratio of blacklist-region reads less than 0.05 were retained for further analysis. Putative doublets identified by ArchR v0.9.3 (***71***) and non-Treg, non-Tconv contaminant cells were also removed. We used the latent semantic indexing approach as previously described (*59, 72*). Binarized count matrices were normalized using the term frequency-inverse document frequency (TF-IDF) transformation and reduced in dimensionality by singular value decomposition (SVD). As the first SVD component was highly correlated with sequencing depth, components 2-30 were used to create a two-dimensional UMAP visualization. A gene accessibility score was determined from Treg scATAC-seq data with ArchR v0.9.3, using an exponentially weighted function that accounts for the activity of distal OCRs in a distance-dependent manner (*71*) and computes a single measure of a gene’s accessibility. A gene-gene correlation was next determined from the log-normalized gene scores and visualized using a force-directed network (edge-weighted using correlation coefficients). The network was filtered to drop weak correlations (-0.2 > and < 0.2). Network clustering was performed using correlation coefficients (edge attribute) and FoxP3 EPL intensities (node attribute). Network visualization and clustering were performed using Cytoscape v3.8.0 (*73*).

### CAGE data

Treg CAGE data were downloaded from the FANTOM repository (*74*).

### H3K27Ac, FoxP3 and Yy1 HiChIP data processing and enhancer-promoter looping

HiChIP paired-end reads were aligned to the mm10 genome using the HiC-Pro pipeline (*75*). Default settings were used for removal of duplicated reads, assignment to MboI restriction fragments and filtering for valid interactions. To assess enhancer-promoter loops (EPLs) in HiChIP data, we computed +/-250 kb contacts from the gene TSS for 9,058 expressed genes in Tregs and Tconv using the hicPlotViewpoint (*76*) function at a resolution of 5 kb. To assess a background proximity ligation frequency for each HiChIP experiment, we first generated a ligation probability background by sampling 1000 randomly selected intragenic regions more than 10 kb from the promoter and computing the relative counts using a window of +/-250 kb in each HiChIP experiment. For each respective distance position, a Z-statistic was computed using the HiChIP background estimation and adjusted p-values were calculated to assess statistical significance of the contact. This procedure removed from 50-80% of noisy and biased chromatin interactions. Significant EPLs were considered if demonstrated enrichment across replicates, passing an FDR of 5% for H3K27Ac EPLs (n=115,538) and 10% for Yy1 (12,426 EPLs), and enriched with respect to the negative control IgG HiChIP. In a similar manner, FoxP3 EPLs were also further analyzed using using a reproducible set of overlapping 13,982 FoxP3 ChIP-seq peaks (*32, 41*) as anchors. All FoxP3 EPLs (10% FDR) were supported by FoxP3 ChIP-seq binding for at least one end of the FoxP3 loop. 1D H3K27Ac HiChIP analysis was performed using all reads (including re-ligation and dangling-ends, but excluding extrachromosomal contacts) and converted into tabular format (bedpe, using cLoops2 (*77*)) for downstream analysis. For each H3K27Ac EPL, 1D HiChIP signal was established for the both TSS and for distal anchor, and these summed. Per-gene 1D H3K27Ac cumulative signal was calculated by summing 1D signal corresponding to all respective EPLs for each gene. Compartment analysis of WT-GFP and KO-GFP HiChIP was performed at 250 kb resolution using the eigenvector function on KR normalized maps (*78*).

### HiChIP experimental metrics

| <u>Sample</u> | <u>Total pairs processed</u> | <u>Unique pairs aligned</u> | <u>Cis intrachromosomal %</u> | <u>Pearson (Inter-replicate)</u> |
| --- | --- | --- | --- | --- |
| <u>H3K27Ac Treg R1</u> | <u>172312254</u> | <u>68%</u> | <u>72%</u> | <u>0.85</u> |
| <u>H3K27Ac Treg R2</u> | <u>140199897</u> | <u>68%</u> | <u>72%</u> |  |
| <u>H3K27Ac Tcon R1</u> | <u>133066435</u> | <u>69%</u> | <u>72%</u> | <u>0.84</u> |
| <u>H3K27Ac Tcon R2</u> | <u>114716225</u> | <u>70%</u> | <u>72%</u> |  |
| <u>Foxp3 Treg R1</u> | <u>53374194</u> | <u>63%</u> | <u>69%</u> | <u>0.82</u> |
| <u>Foxp3 Treg R2</u> | <u>51874296</u> | <u>62%</u> | <u>69%</u> |  |
| <u>Yy1 Treg R1</u> | <u>37509324</u> | <u>61%</u> | <u>68%</u> | <u>0.8</u> |
| <u>Yy1 Treg R2</u> | <u>30881902</u> | <u>62%</u> | <u>68%</u> |  |
| <u>IgG</u> | <u>8328069</u> | <u>59%</u> | <u>69%</u> |  |
| <u>H3K27Ac WT GFP R1</u> | <u>121381491</u> | <u>58%</u> | <u>70%</u> | <u>0.85</u> |
| <u>H3K27Ac WT GFP R2</u> | <u>139482053</u> | <u>53%</u> | <u>69%</u> |  |
| <u>H3K27Ac KO GFP R1</u> | <u>99594436</u> | <u>63%</u> | <u>72%</u> | <u>0.8</u> |
| <u>H3K27Ac KO GFP R2</u> | <u>108433482</u> | <u>60%</u> | <u>72%</u> |  |

### Cumulative EPL intensities determined for genes

To measure the overall effect of the enhancer-promoter structure for a given gene, we calculated a single cumulative value by summing the respective HiChIP EPL intensities (H3K27Ac, FoxP3, Yy1). Additionally, the degree in connectivity was calculated by summing the total number of EPLs per gene and compared both with genome-wide expression and binned mRNA quantiles.

### Super interactive promoter analysis

To identify Super interactive promoters from Treg and Tconv H3K27Ac or FoxP3 HiChIP, we followed a similar strategy employed by (*48, 79*). Briefly, for each gene we summed the log10 FDR for respective EPLs to derive a single cumulative score per gene. Plots of the ranked cumulative interaction scores in each cell type and condition were plotted. Significant SIPs were determined based on the point of inflection within the curve (slope is equal to 1).

### Differential H3K27Ac EPL analysis

Differential H3K27Ac EPLs were identified using biological replicate EPL counts of Treg and Tconv EPLs (P.adj < 0.05, Fold change > 2) determined using diffloop (*80*). Cumulative H3K27Ac EPL differences and comparisons with gene expression between T cells were mapped for all genes and specifically highlighted for the up- and down- regulated Treg signature program (*39*).

### Yy1 and FoxP3 comparative analysis

Yy1 EPL and cumulative intensities were compared with FoxP3 respectively to identify TF-specific interactions (2-Fold, P<0.05). Gene ontology analysis was performed for genes with differences between FoxP3 and Yy1 cumulative EPL intensities (2-Fold) using Metascape (*81*).

### Deriving a FoxP3-dependent signature from RNA-seq data

Data from GSE154680 (*52*) were downloaded. Raw read counts tables were normalized by median of ratios method with DESeq2 (*82*) and converted to GCT and CLS formats. Genes with a minimum average read count of 35 in at least one group and a coefficient of variation intra-replicate less than 0.30-0.45 were retained. An uncorrected t-test was used to compute differential gene expression between the different groups from the normalized read counts dataset. The FoxP3 dependent gene signature list was computed by merging the differential expressed genes from the aTreg and rTreg datasets comparing Foxp3.GFP.DTR WT versus Foxp3.GFP KO and Fold change >2 or <0.5 and p-value < 0.01.

### Promoter, enhancer, and dual modes in FoxP3 looping

FoxP3 ChIP-seq binding data was used to parse FoxP3 EPLs into those with FoxP3 occupancy at promoter, enhancer or both (dual mode) ends of enhancer-promoter loops. Genomic heatmaps (69) were generated for each group either centered on FoxP3 bound positions in FoxP3 EPLs or centered on the strongest Treg ATAC-seq OCR for FoxP3-negative sides of the EPL. Gene ontology analysis of FoxP3 modes were generated using Metascape (*81*). *De novo* motif analysis was performed using HOMER for FoxP3 bound positions of promoter and enhancer EPL modalities. Significant motifs were determined using as background comparisons using equal numbers (10, 000) in TSS and non-TSS OCRs. Motif significance was represented for promoter and enhancer FoxP3 modes using Circos plots (*83*).

### *Il2* and *Il2ra* centric loop analysis

To resolve higher-resolution sets of chromatin loops, we first defined potential anchors from OCRs with accessibility in Tconv or Tregs (297,076 OCRs). HiChIP loops were defined from our OCR anchor set using Hichipper (*84*) (-peak-pad 250 bp), which adopts a restriction fragment bias-aware approach for preprocessing of HiChIP data to call loops. Briefly, anchors were padded 250 base pairs in either direction and were considered if they overlapped MboI motifs. Padding anchors boosts the number of paired-end tags that can be mapped to loops respectively. We removed self-ligation loops and filtered loops that may be biased due to proximity (< 3kb) or low paired-end tag counts using the Mango (*85*) correction step (stage 5). This correction aggregates paired-end tag counts across all samples and retains significant loops (FDR < 0.01). A union set of loops across all data was generated; 1) loops with a minimum of 2 counts in both biological replicates, 2) supported in at least 1 sample across all HiChIP experiments, and 3) greater than 2-Fold compared to IgG HiChIP controls. We mapped the hichipper defined loops based on overlapping HiChIP EPLs for FoxP3, H3K27Ac and Yy1 centering on the *Il2ra* and *Il2* locus.

### Data visualization

Analysis was performed using R-3.5.2 with all plots generated using ggplot2 (https://ggplot2.tidyverse.org) (*86*). Statistical tests are described in their respective sections. Browser and genome views of ATAC-seq and ChIP-seq data were generated using UCSC (*87*) and IGV (*88*). EPL arc plots were generated using DNARchitect (*89*). HiChIP heatmaps were generated using HiC-Pro by merging valid pairs of replicates and processed as .hic files using the hicpro2juicebox function (*75*) and visualized using Juicebox tools (*78*).

## ACKNOWLEDGEMENTS

We thank: Drs. Caleb Lareau, Rick Young, and Nicole El-Ali for insightful discussions; K. Hattori, C. Araneo, K. Seddu, and the HMS Biopolymers team, for help with mice, cell sorting and sequencing. This work was funded by grants from the NIH to CB&DM (AI116834, AI150686), JL was supported by an INSERM Poste d’Accueil and an Arthur Sachs scholarship, RR by NIH supplement AI116834-03S1, and KC by NIGMS-T32GM007753 and a Harvard Stem Cell Institute MD/PhD Training Fellowship.

## Data and materials availability

All data and materials are available upon request from the corresponding author. All data needed to evaluate the conclusions in the paper are present in the paper.

## SUPPLEMENTAL FIGURE LEGENDS

**Figure S1.**
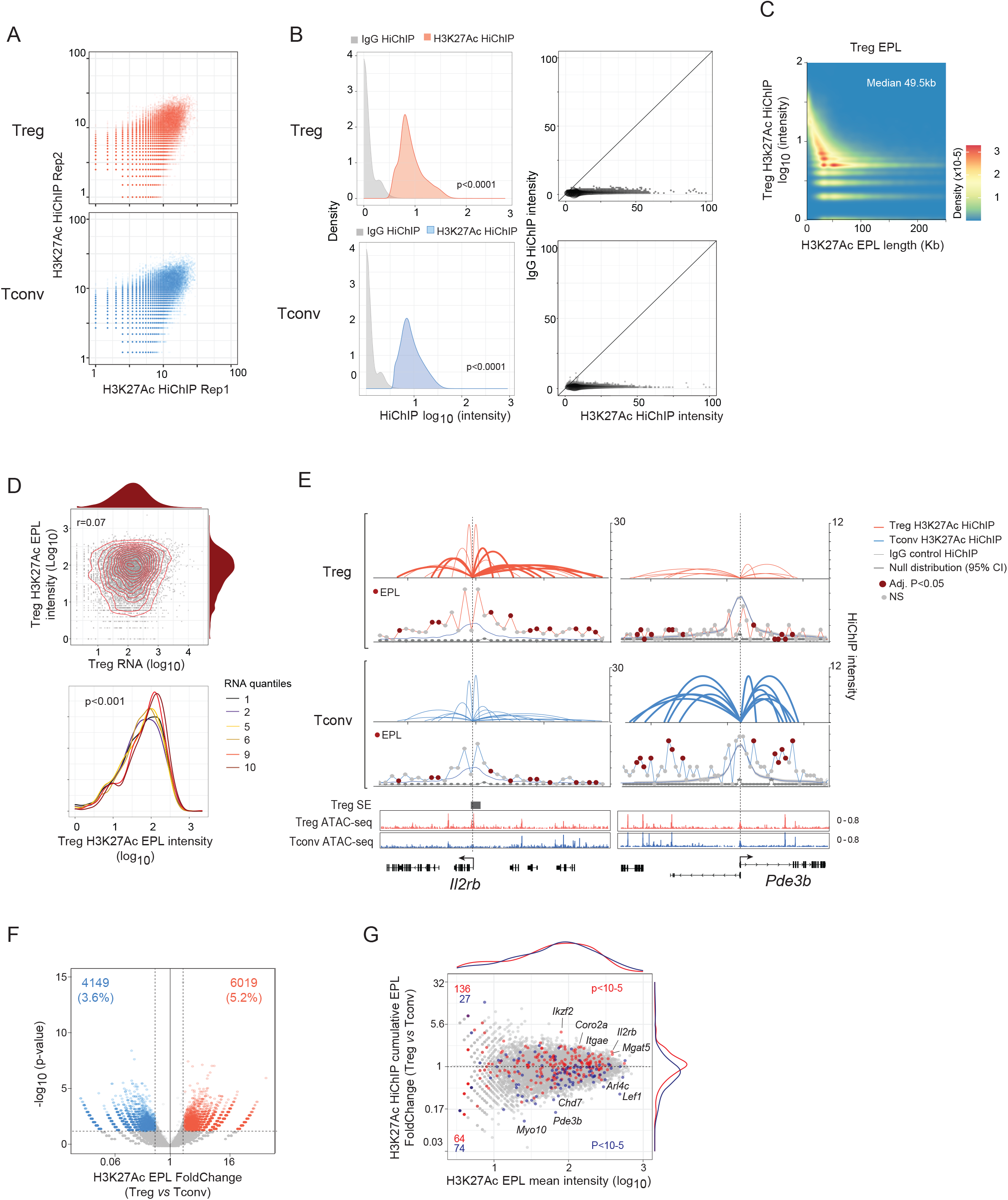
Mapping the H3K27Ac enhancer-promoter architecture of conventional and regulatory T cells. A. H3K27Ac HiChIP EPL biological replicate correlations for Treg (top, Pearson=0.85) and Tconv (bottom, Pearson=0.84). B. Quality control analysis of H3K27Ac Treg (top) and Tconv (bottom) HiChIP using IgG HiChIP control. Density (left) and scatter (right) plots compare H3K27Ac and IgG HiChIP intensities among EPLs for each respective cell-type. C. Distribution of H3K27Ac HiChIP EPL intensities (log10) and relative loop size (Kbp) in Tregs, with a median H3K27Ac EPL length of 49.5 kb. D. Genome-wide comparative analysis of cumulative H3K27Ac HiChIP EPL intensities (log10) and mRNA expression (log10) for 9,058 genes (r=0.07, top). Distribution of cumulative H3K27Ac HiChIP EPL intensities (log10) for genes binned as quantiles based on low (10-20), middle (50-60), and high (90-100) expression in Tregs (bottom). E. Vignettes of the T cell-specific enhancer-promoter structures for Treg signature genes *Il2rb* (Up-regulated) and *Pde3b* (Down-regulated). F. Fold change vs. p-value (volcano) plot comparing H3K27Ac HiChIP EPLs in conventional and regulatory T cells. EPLs with differential intensity (adj.P<0.05, FC>2) are highlighted with numbers shown. G. Fold change vs. cumulative H3K27Ac HiChIP EPL intensities, highlighting the Treg Up- and Down- gene signature (red and blue, respectively). Significance in the EPL intensity deviations were determined using a one-way Chi-square test in Up (P<10^-5^) and Down (P<10^-5^) gene subsets respectively.

**Figure S2.**
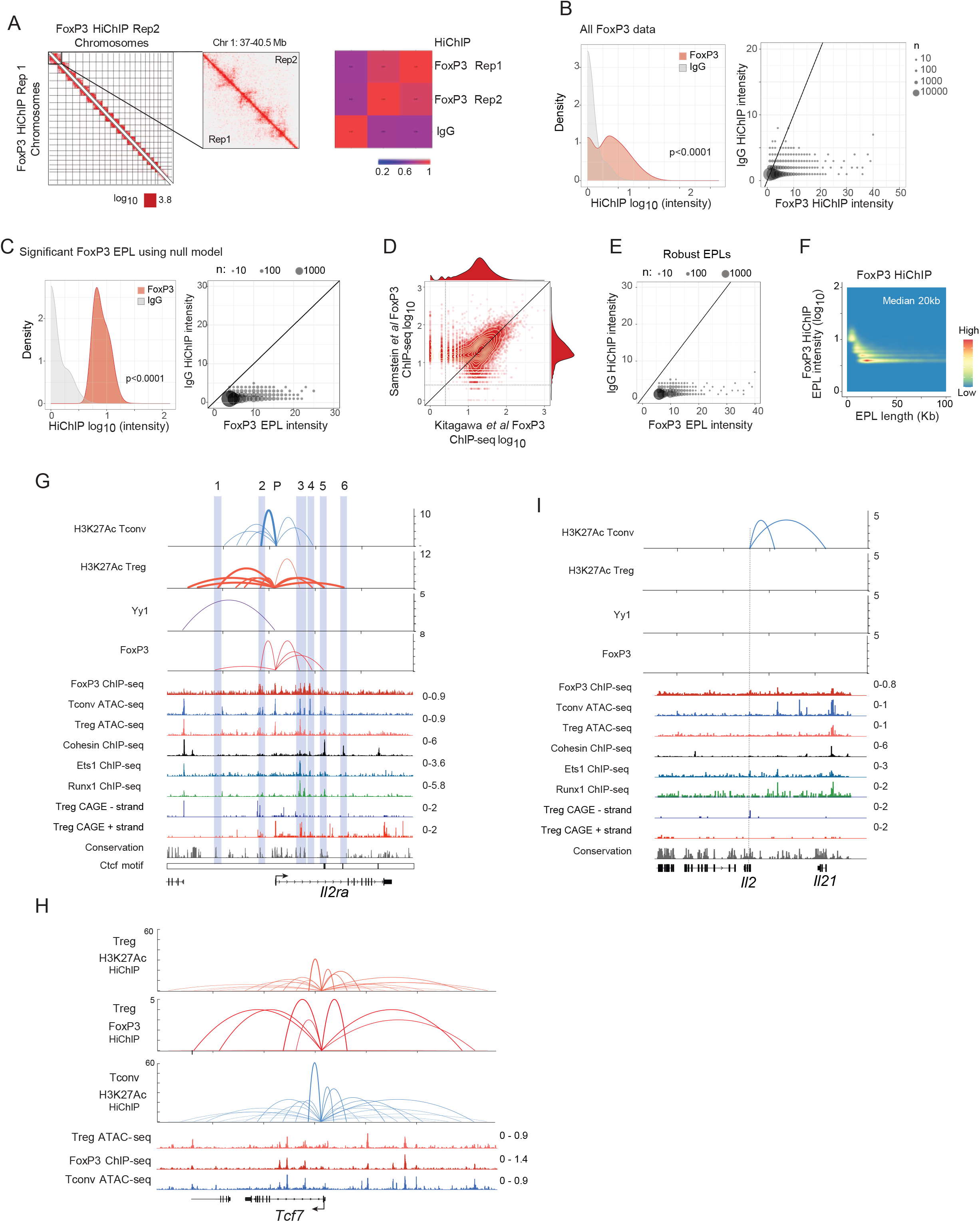
Determining Treg FoxP3 enhancer-promoter loops using HiChIP. A. Genome-wide chromosomal view of FoxP3 HiChIP biological replicates with a focused inset on chr1 and the correlations (Pearson) for FoxP3 HiChIP replicates and IgG control HiChIP (right). B. Density (left) and scatter (right) plot comparisons of FoxP3 HiChIP and IgG intensities based on all unfiltered EPLs. C. Density (left) and scatter (right) plot comparisons of FoxP3 HiChIP and IgG intensities for high-confidence EPLs following analysis using the proximity-biased null distribution. D. Comparison of FoxP3 ChIP-seq datasets from (*41*) (x-axis, log10) and (*32*) (y-axis, log10). Lines (>3 ChIP-seq counts) emphasize the FoxP3 ChIP-seq peaks concordant in both datasets. E. Scatter plot comparison of FoxP3 (x-axis) and IgG (y-axis) HiChIP intensities for high-confidence and FoxP3 ChIP-seq supported FoxP3 HiChIP EPLs. F. Distribution of FoxP3 HiChIP EPL intensities (log10) and length of FoxP3 EPLs (Kb), with a median size 20 Kb. G. Genomic view of H3K27Ac (Tconv and Treg), FoxP3 and Yy1 loops for the *Il2ra* locus. Highlighted regions mark positions of FoxP3-specific HiChIP loops. TF ChIP-seq and CAGE data below are profiled in Tregs. H. Genomic view of H3K27Ac (Tconv and Treg) and Treg FoxP3 loops for the *Tcf7* locus. I. Genomic view of H3K27Ac (Tconv and Treg) and FoxP3 and Yy1 loops for the *Il2* locus. TF ChIP-seq and CAGE data below are profiled in Tregs.

**Figure S3.**
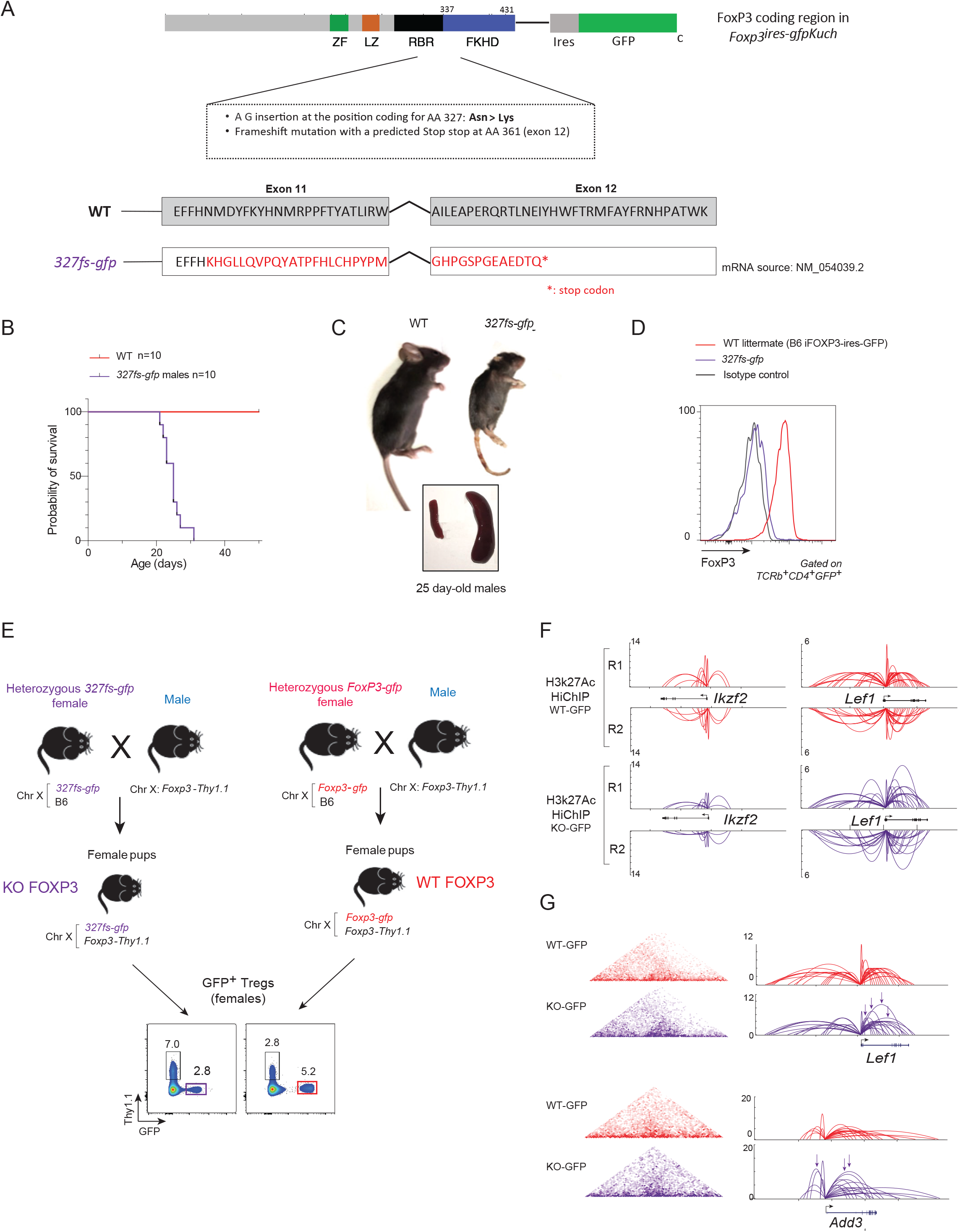
Generation and validation of FoxP3-deficient mice. A. Position and consequence of the mutation in FoxP3-deficient *Foxp3fs327-gfp/Doi* allele generated by CRISPR mutagenesis of the *Foxp3-gfp/Kuch* reporter allele (*54*). RBR: Runx-binding region; FKHD: Forkhead domain B. Survival curve of *B6.Foxp3.fs327-gfp* mice. C. Images of WT and *B6.Foxp3.fs327-gfp* mice.and their spleens D. FoxP3 expression detected by intracellular staining and flow cytometry in CD4+ spleen Treg and Treg-like cells. Note the very slight shift in *Foxp3.fs327-gfp* relative to unstained control. E. Breeding strategy to obtain comparable Treg-like cells in which the *Foxp3.fs327*WT or *Foxp3-gfp/Kuch* alleles are active, in heterozygous females protected from inflammation by the normal *Foxp3-Thy1.1* reporter allele. F. Reproducibility in H3K27Ac HiChIP EPL intensities between biological replicates for WT-GFP (top) and KO-GFP (bottom). The left panels show H3K27Ac EPLs for the *Ikzf2* locus and those for *Lef1* on the right. G. WT-GFP (red) and KO-GFP (purple) H3K27Ac HiChIP intensity maps (left) and EPLs (right) of FoxP3 targets *Lef1* and *Add3*. Arrows indicate EPLs FC>1.5 and P<0.05).

**Figure S4.**
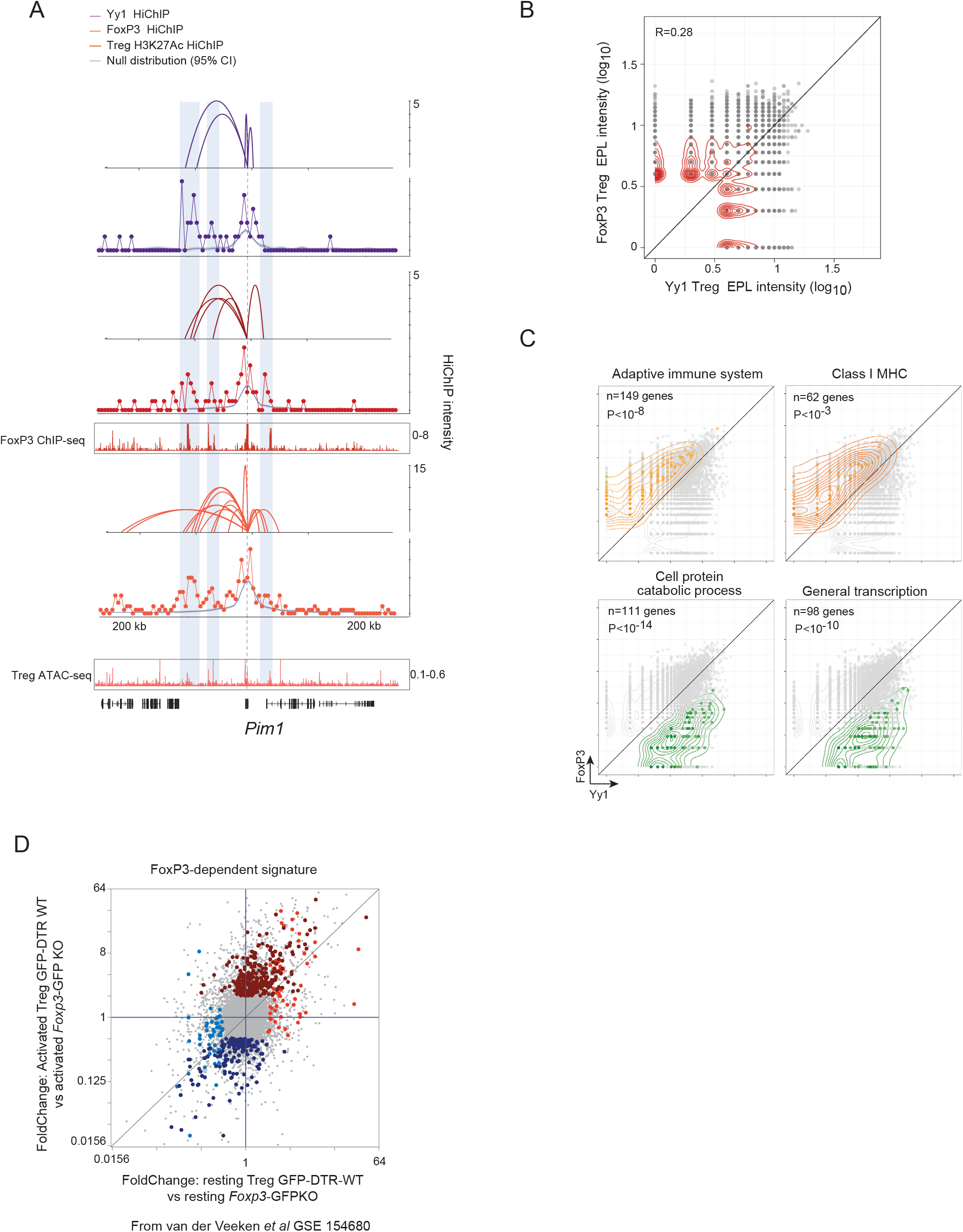
Transcriptional specificities detailed by enhancer-promoter topologies of FoxP3 and Yy1 in regulatory T cells. A. Treg Yy1 (top), Treg FoxP3 (middle), and Treg H3K27Ac (bottom) enhancer-promoter loops for the *Pim1* locus. Highlighted areas show differences of Yy1 and FoxP3 EPLs in Tregs. B. Scatter plot comparing Yy1 (x-axis, log_10_) and FoxP3 (y-axis, log_10_) HiChIP EPL intensities (Pearson corr. 0.28). C. Gene-level cumulative EPL intensities comparing Yy1 (x-axis, log_10_) and FoxP3 (y-axis, log_10_). Gene ontology enrichments are shown for genes enriched (>2 Fold) in FoxP3 HiChIP (top, orange) or Yy1 HiChIP (bottom, green) EPL intensities. D. FoxP3-dependent signature genes (n=576) (*52*) shown for comparisons within resting (x-axis, Treg GFP-DTR-WT vs Foxp3-GFPKO) and activated Tregs (y-axis, Treg GFP-DTR-WT vs Foxp3-GFPKO). Genes colored red are upregulated, those in blue are downregulated FoxP3-dependent genes.

**Figure S5.**
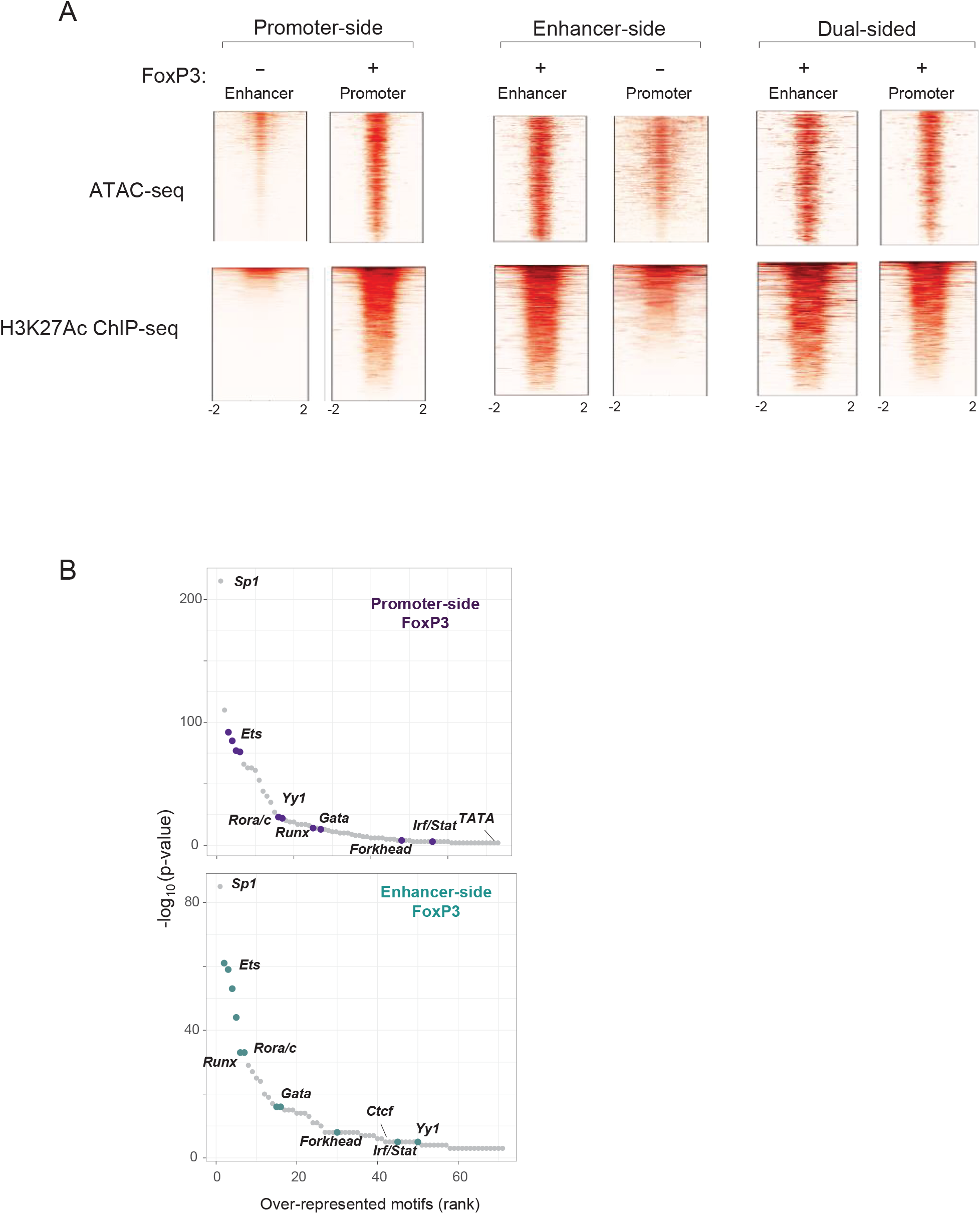
FoxP3 EPL modalities reveal differences in TF motif enrichments. A. Heatmaps of FoxP3 HiChIP EPLs based on promoter, enhancer and dual modes. Treg ATAC-seq (top) and H3K27Ac ChIP-seq (bottom) signals are shown for each mode. Enhancer sides are positioned to the left and promoter to the right for each group respectively. B. Ranking of TF motif enrichments (-log_10_ P-value) of promoter-side (top) and enhancer- side (bottom) FoxP3 EPLs.

## SUPPLEMENTAL TABLE LEGENDS

Table S1 T cell EPLs. Tconv and Treg H3K27Ac EPLs, Treg FoxP3 EPLs and Yy1 EPLs are listed per gene for genomic position and intensity for each respective HiChIP experiment. NA’s
indicate absence of EPL for the respective experiment.

Table S2 Cumulative EPL intensity and T cell Super Interactive Promoters (SIP). EPL cumulative intensities for H3K27Ac, FoxP3 and Yy1 experiments. Significant H3K27Ac SIPs are listed for Tconvs and Tregs.

Table S3 FoxP3-dependent coregulated network. Gene and respective coordinate information for the genes selected in fig. S3D and displayed in Fig. 4. Cluster designation and FoxP3 EPL intensities are included. RNA expression values, fold change and p-values are included for Treg RNA-seq datasets from GSE154680.

Table S4 Gene ontology enrichments of FoxP3 EPLs. Top gene ontology enrichments and respective genes are listed for each FoxP3 EPL class.

## Supplementary Note

### Indirect regulation is not FoxP3’s dominant mode of action

T regulatory (Treg) cells characterized by the transcription factor FoxP3 are central elements of peripheral immune tolerance. These CD4+ T cells act as dominant negative regulators of many facets of the immune system, controlling immune responses and enforcing peripheral tolerance to self-antigens, symbiotic commensals or fetal antigens (6). Treg cells are also involved in controlling non-immunologic consequences of inflammation in several tissular locations (7). The Treg lineage is determined by the transcription factor (TF) FoxP3, which partly shapes its transcriptome. This dependence is only partial, as Treg-like cells (“Treg wannabes”) are observed even in the complete absence of FoxP3 [(8) and refs therein]. Accordingly, only a portion of the Treg-specific transcriptome is FoxP3-dependent [(9) and refs therein]. But, even if not the master regulator it was once portrayed as, FoxP3 does remain a determining player in shaping Treg identity and function, its importance illustrated by the devastating autoimmune pathology in FoxP3-deficient mice and humans.

There has been considerable interest in the mechanics of FoxP3’s control on its target genes, from structural and mechanistic standpoints, but the question remains unresolved. It has been generally assumed that FoxP3 follows the traditional model of TF activity, that it activates or represses target genes by binding their cis-regulatory elements, synergizing with other TFs at those locations (Fig. 1A). A number of reports supports this view (1,10–13). A new model recently proposed by van der Veeken et al (5) posits that FoxP3 shapes Treg identity indirectly, by tuning the tctivity of intermediary TFs, in particular TCF-1 (Fig. 1B). This ”indirect model” bears some relation to the “TF theft” recently proposed for PU.1 (14) - in both cases, action through reducing the availability of an intermediate effector TF. This conclusion was reached through a series of carefully and elegantly strung arguments, which proceed as follows (1) The first step involved defining the scope of FoxP3-dependent transcripts. From there, it was shown that (2) the majority of FoxP3-binding sites have no transcriptional consequence; (3) conversely, there is no enrichment in FoxP3-binding among FoxP3-dependent loci. (4) From these two tenets derived the inference that FoxP3-dependent changes must be mediated indirectly, through the activity of one or more TFs whose expression levels are affected by FoxP3. (5) Validating the hypothesis, experimentally reducing the levels of one of the proposed intermediate TFs, TCF-1, recapitulated FoxP3-dependent effects on chromatin.

**Figure 1a.**
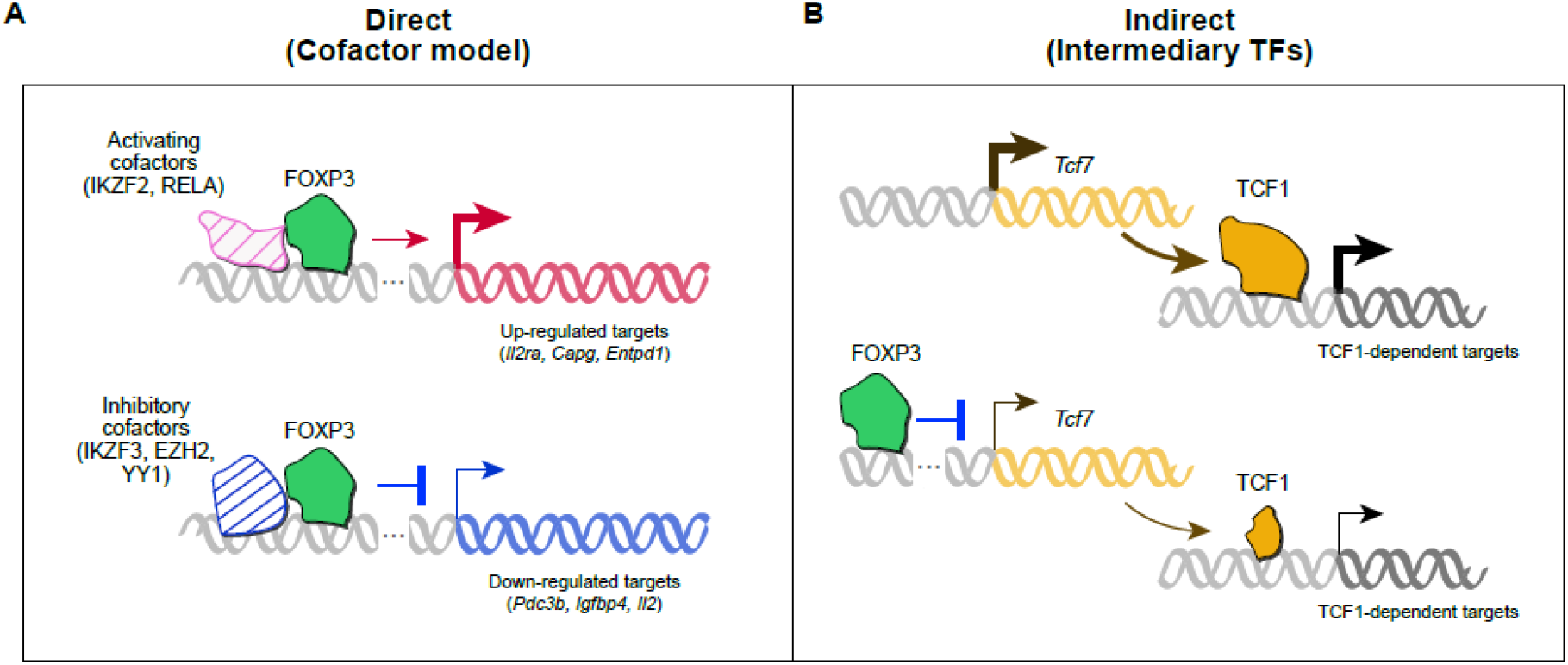
Models of FoxP3 mode of activity (generic for any transcription factor) Different modes of action have been proposed to account for FoxP3’s transcriptional regulation in Treg cells, and how it can activate the expression of some transcripts while repressing others **A.** The generic direct mode of transcription factor activity (here “Cofactor model”) is the one most generally considered for transcription factors: binding to a cognate recognition sequence (close to the promoter region of a target gene, or at a distal enhancer) allows the interaction with another transcription regulator in the vicinity (where this co-binding may actually be facilitated by either partner). The nature of the co-factor is what conditions the resulting activating or inhibitory activity. Mechanistically, there may be different modalities to activation or repression: enzymatic activities that modify Pol-II, nucleosomes or ancillary actors in transcriptional regulation; further recruitment of additional complexes; modification of enhancer-promoter loops in nuclear DNA. Note that the relationship between FoxP3 and a particular target gene need not be mono-deterministic. It may be *probabilistic* (not operational in every single cell within a given Treg subtype); or the co-factor recruited by FoxP3 at a given site may vary according to post-translational modifications of the FoxP3 protein (4). **B.** Indirect regulation, where “intermediate” TFs (here TCF1) positively regulate target genes, and are themselves repressed by FoxP3. Some transcripts might thus appear dependent on FoxP3, but wouldn’t be direct targets. It was recently proposed that this is FoxP3 principal mode of action (5).

However, we believe that the string of arguments does not make a watertight case for the model stated unequivocally in the paper’s title. It is certainly plausible that some of FoxP3’s effects occur indirectly, indeed likely given that some TFs are indeed FoxP3 targets. But we find that this mechanism is not a major component of FoxP3’s contribution to Treg cell determinism and transcriptome, in particular when considering loci that are essential to Treg cell physiology, like *Il2ra*. We will point out, in sequence, some ambiguities or alternative explanations to the logical path mentioned above.

#### 1. Definition of FoxP3-dependent genes

Defining the scope of FoxP3-dependent genes is essential to any discussion of FoxP3’s mechanism of action. The paper uses an elegant approach, comparing FoxP3-inactivating vs non-inactivating GFP insertions (equivalent to WT and KO, but both reported by *Foxp3*-driven GFP for easy identification and sorting). This in heterozygous females, where the WT Tregs ensure immune homeostasis. In principle, this FoxP3-dependent list will include some genes directly regulated by FoxP3, and others under indirect control. This definition is probably the most rigorous that can be proposed (and we use it in our own work (15)], but there are some caveats, which suggests that the scope of FoxP3-dependent genes may have been over- estimated.

- This definition rests on the assumption that the Treg-like “wannabes” present in FoxP3-deficient mice are normal Tregs that solely lack FoxP3 and FoxP3-dependent transcription. This assumption may be uncertain: the absence of FoxP3 may be compensated by activation of complementing TFs, cells may be unstable and turnover more rapidly. Prosaically, wannabes may be “oddballs”, with transcriptome alterations that would be mistakenly included as FoxP3-dependent. Along the same lines, our recent single-cell RNAseq analysis of FoxP3-deficient mice shows that Treg wannabes are a mixed population, possibly even including some activated conventional T cells (9).

- In addition, there is only limited overlap with the list of FoxP3-dependent genes determined by direct transduction of FoxP3 into CD4+ T cells (1) (21/580, 3.6%). Certainly, this alternative definition is itself open to potential artefacts (FoxP3 transduction does not fully recapitulate Treg identity, FoxP3 transduction into CD4+ T cells requires strong cell activation, which may lead to confounders, and the usual issues with enforced TF expression). But one might have wished better concordance.

#### 2. The majority of FoxP3-binding sites have no obvious transcriptional consequence

No contest here. Tracing back to the original analyses of FoxP3 binding genome- wide (2, 3), it has been recognized that FoxP3 can bind many locations in the genome that are nowhere near any FoxP3-depdendent gene. Results from our recent analysis of FoxP3 binding to enhancer-promoter loops (15) further indicate that this “irrelevant binding” occurs at actual enhancer-promoter loops, not in inert genomic locations. Widespread TF binding beyond loci where they are actually active is not specific to FoxP3, and a common occurrence (16, 17). This “irrelevant binding” has been proposed to function as a TF reservoir, a buffer, or may simply be the unavoidable consequence of mass action law and of the low complexity of TF-binding motifs and their wide distribution across DNA. In the cofactor model, irrelevant binding corresponds to locations where FoxP3 does not find appropriate cofactors or modifiers in order to be active. Thus, this disconnect is not necessarily a strong argument for indirect regulation.

#### 3. There is no enrichment in FoxP3-bound genes among FoxP3-dependent loci

This conclusion, the flip side of the previous, mostly derived from the data shown in Fig. 4B/C of ref (5), but we believe that there may have been a misinterpretation. There **is** an enrichment of FoxP3-bound loci among FoxP3-dependent genes (e.g. calculating from Fig. 4C numbers: within FoxP3-bound 146/7029=2.0% are up-regulated in FoxP3 WT/KO; for all genes, 302/45375= 0.6%, and thus a 3.1-fold enrichment). The same enrichment is seen in resting and activated Treg cells. Similarly, using the identification of FoxP3-dependent genes from transfection data (1), we again observe an enrichment of FoxP3-bound genes within FoxP3-dependent genes, especially in the up-regulated contingent.

#### 4. FoxP3-dependent changes must be indirect via tuning of other TFs

“Thus, the pervasive Foxp3-dependent chromatin accessibility and gene expression changes at non-bound sites must be mediated indirectly through the activity of one or more Foxp3-dependent trans-regulatory factors”.

Even if points 2 and 3 were given, this inference would be a strong logical jump, as other interpretations are also plausible to explain the discordance between changes in gene expression and apparent FoxP3 binding, without involving indirect effects through other TFs. For one, because this discordance may be less important than it seems. It is well known that techniques based on chromatin immunoprecipitation (ChIP-seq, CUT&RUN, etc) do not necessarily report on the entire range of TF binding modalities. Aside from technical artefacts (18) which are not in question here (results from the Rudensky and Sakaguchi labs are very concordant (2, 3)], chromatin immunoprecipitation favors stable interactions over dynamic ones (e.g. much more effective for histone modifications or architectural TFs like CTCF than for DNA-binding TFs); some TFs prove refractory to ChIP altogether, even with good affinity antibodies (19, 20). Recent high-resolution single molecule occupancy studies show that TF cooperativity is required for nucleosome displacement by mechanisms that may not be captured by conventional ChIP (21). Thus, the discordance between FoxP3 binding assessed by immunoprecipitation and functional activity may reflect the limitations of ChIP. FoxP3 function might involve short-lived interactions, and some of the stable landing sites detected by ChIP might even be red herrings.

**Table 1.**
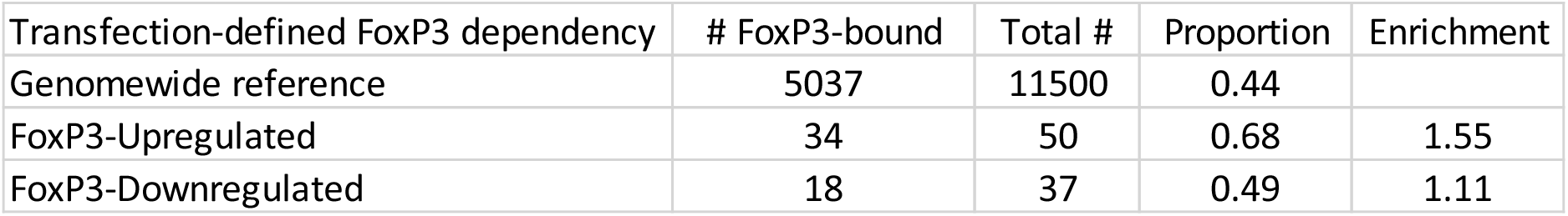
Enrichment of FoxP3-bound genes within those functionally activated by FoxP3. FoxP3-up and -downregulated genes, as defined by transfection into CD4+ T cells (1), cross-matched with FoxP3-binding genes (robustly defined by confirmatory discovery in FoxP3 ChIP-seq from Sakaguchi and Rudensky laboratories (2, 3).

| Transfection-defined FoxP3 dependency | # FoxP3-bound | Total # | Proportion | Enrichment |
| --- | --- | --- | --- | --- |
| Genomewide reference | 5037 | 11500 | 0.44 |  |
| FoxP3-Upregulated | 34 | 50 | 0.68 | 1.55 |
| FoxP3-Downregulated | 18 | 37 | 0.49 | 1.11 |

Further, some directly-acting TFs may not require cognate binding at all. Either because they are recruited to pre-existing complexes (Aire is in this category), and regulate transcription through enzymatic modification (e.g. manipulating the local histone code). Thus, observing no ChIPseq or CUT&RUN signal does not negate a direct activity of a TF in a genomic location.

#### 5. Lowering of TCF-1 levels recapitulate FoxP3-dependent changes

This elegant attempt at confirming the indirect model is reported in Figs 5 and 6 of ref (5): loci apparently FoxP3-dependent, but in fact controlled by an intermediate TF, would be sensitive to experimentally tuning the levels of this intermediate. However, this experiment doesn’t meet its goal.

- Most directly, the statement “*Tcf7 hypomorph cells did not exhibit strong differential expression of FoxP3-dependent genes*” basically negates the whole design. It is certainly possible that redundancy between TCF-1 and LEF1 (13) explains the lack of effect, but this key test fails to provide a proof-of-principle of the model.

- In fact, the opposite effect is observed in fig. S5C. The indirect-control model (FoxP3 represses TCF-1, and TCF-1-activated transcripts thus appear repressed by FoxP3) predicts that genetically lowering TCF-1 levels would lower the expression of FoxP3-downregulated genes. The opposite happens: the majority of genes down- regulated by FoxP3 are *up-regulated* in TCF-1-underexpressing cells. And *vice versa,* genes up-regulated by FoxP3 are repressed in TCF1 hypomorph Treg cells (fig. S5C of (5). Beyond the mind-bending double-negatives, this outcome is more in line with the notion that TCF-1 may actually cooperate *positively* with FoxP3 to regulate a number or FoxP3-dependent targets.
- At quick glance, Fig. 5G of ref (5) could be interpreted as establishing a connection between TCF-1 binding and FoxP3-dependent expression. It is misleading, however, as the genesets whose expression is skewed were selected on the basis of TCF-1 binding AND of differential ATAC-seq in FoxP3-proficient and –deficient cells. An essential control is missing (genes with the same shift in ATAC signal, irrespective of TCF-1 binding).

#### Arguments in favor of the “cofactor model” in the van der Veeken results

On the other hand, several of the results presented in the data-rich ref (5) could be interpreted to support the cofactor model of direct control by FoxP3.

- Ref (5) reports an elegant identification of DNA motifs that drive epigenetic features, leveraging naturally occurring genetic variation between distant mouse species, by correlating variation in TF-binding motifs with local intensity of epigenetic signals (22, 23). Fig. 3E highlights TF-binding motifs whose match with optimal binding coincides with increased binding of FoxP3, detected by ChIPseq or CUT&RUN (Fig. 3E). Forkhead-binding elements show up as positively correlated with FoxP3 binding (a good positive control) but motifs for other TF families do so as well (Ets, bZIP, ZF), indicating that stable FoxP3 binding is positively facilitated by its cofactors.
- Fig. 5D displays differential chromatin accessibility, at TCF-1 binding sites, in WT or FoxP3-deficient Treg wannabes. Some loci bound by TCF-1 alone show decreased accessibility when FoxP3 is present in the cell, plausibly interpreted as showing that FoxP3 lowers TCF-1 expression, and hence accessibility at TCF-1 dependent sites (5). But this drop is surprisingly not observed for genes bound by *both* TCF-1 and FoxP3, where the same logic would theoretically apply. This paradox suggests that FoxP3 protein may compensate for the TCF-1 deficiency by restoring chromatin accessibility at TCF-1- dependent loci. Hence, even though FoxP3 antagonizes TCF-1 by repressing its expression, it may paradoxically cooperate with it at the functional level. This situation is reminiscent of a paradoxical cooperativity that we described earlier between FoxP3 and Satb1 or Lef1: even though these TFs are under-expressed in Treg cells relative to conventional CD4+ T cells, they actually cooperate with FoxP3 in locking in Treg identity (13). If one also considers RORγ, it may be an emerging theme that antagonism at the level of expression accompanies cooperativity at the level of function.

In summary, while the indirect mode of regulation by FoxP3 certainly occur, it does not dominate FoxP3’s mode of action as had been suggested, and co-exists with the more variegated regulatory makeups that the combinatorial features of the cofactor model allows.

## Notes

### Competing Interest Statement

The authors have declared no competing interest.

